# ASCL1 drives induction of a transitory cell state required for repair of the injured neonatal brain

**DOI:** 10.1101/2020.11.03.366872

**Authors:** N. Sumru Bayin, Dogukan Mizrak, Daniel N. Stephen, Zhimin Lao, Peter A. Sims, Alexandra L. Joyner

## Abstract

The underlying molecular foundation of neural progenitor diversity and plasticity is critical for understanding repair processes. The neonatal cerebellum with multiple progenitor populations has high regenerative potential. Upon ablation of cerebellar granule cell progenitors at birth, a normally gliogenic Nestin-expressing progenitor (NEP) in the Bergmann glia (Bg) layer (BgL-NEPs) undergoes adaptive reprograming to restore granule neuron production while a white matter NEP (WM-NEPs) reduces interneuron production. However, the cellular states and genes regulating the NEP fate switch are not known. Here using scRNA-seq and fate-mapping, we defined the molecular subtypes of NEPs and their lineages under homeostasis and repair. Five NEP populations comprising two molecular subtypes, *Hopx*-expressing gliogenic- and *Ascl1*-expressing neurogenic-NEPs were identified in both states. Furthermore, in the WM, distinct NEP populations generate interneurons or astrocytes, and amongst gliogenic-NEPs, astrocyte and Bg lineages are molecularly separable. Importantly, we uncovered that after injury a new transitory cellular state arises from *Hopx*-NEPs in the BgL that is defined by initiation of expression of the neurogenic gene *Ascl1*. Moreover, *Ascl1* is required for adaptive reprogramming and the full regenerative capacity of the cerebellum. We thus define new populations of NEPs and identifed the transcription factor responsible for inducing a transitory cell critical for a glial to neural switch *in vivo* following injury.

## Introduction

Progenitor cells must have a multiplicity of subtypes and flexibility in fate choices for complex tissues to be generated and repair of injuries to be effective. Development of single cell approaches has been instrumental in dissecting progenitor diversity and identifying transitory cell states critical to fate decisions (Llorens-Bobadilla et al., 2015; Wang and Navin, 2015). The neonatal cerebellum (CB) represents a valuable system to identify molecular mechanisms that drive plasticity after injury due to its high regenerative potential and diversity of progenitor populations. The CB, a folded hindbrain structure that houses the majority of the neurons in the brain (Azevedo et al., 2009; Herculano-Houzel et al., 2006), is important for motor and higher order cognitive functions (Caligiore et al., 2017; Fatemi et al., 2012; Koziol et al., 2014; Lackey et al., 2018). The CB has prolonged development compared to the rest of the brain as production of a majority of the cells occurs after birth in mammals. The postnatal progenitors of the CB continue proliferating up to 6 months after birth in humans (Rakic and Sidman, 1970) and 2 weeks in mice (Altman and Bayer, 1997). The late development of the CB leads to increased susceptibility to injury around birth and cerebellar hypoplasia is the second leading risk factor of autism spectrum disorders (Wang et al., 2014). Interestingly, the newborn rodent CB can efficiently replenish at least two of its main cell types when they are ablated (Altman and Anderson, 1971; Altman et al., 1969; Bayin et al., 2018; Wojcinski et al., 2017; Wojcinski et al., 2018), and one repair process involves unexpected progenitor plasticity and a glial to neural fate switch after injury. The molecular mechanisms that drive progenitor plasticity *in vivo* however are unknown in the neonatal CB.

After birth, several distinct cerebellar progenitor populations derived from the rhombic lip (RL) or the ventricular zone (VZ) continue to proliferate and generate late-born cells. The rhombic lip-derived *Atoh1*-expressing granule cell progenitors (GCPs) proliferate in the external granular layer (EGL) on the surface of the CB and upon their last cell division, produce excitatory neurons that migrate inwards to form the inner granular layer (IGL) (Machold and Fishell, 2005; Wang et al., 2005; Wingate and Hatten, 1999). A poorly defined group of cerebellar ventricular zone-derived *nestin*-expressing progenitors (NEPs) give rise to astrocytes, inhibitory interneurons (INs) and Bergmann glia (Bg), a specialized polarized glial cell with fibers extending to the cerebellar surface (Cerrato et al., 2018; Fleming et al., 2013; Lee et al., 2005; Milosevic and Goldman, 2004; Parmigiani et al., 2015; Wojcinski et al., 2017). Fate mapping studies have indicated that the lineage propensity of NEPs depends on their position, such that NEPs in the layer that houses the Bg (BgL) or in the prospective white matter (WM) produce Bg or IN, respectively, and that both produce astrocytes (Cerrato et al., 2018; Fleming et al., 2013; Parmigiani et al., 2015; Wojcinski et al., 2017). Furthermore, there are NEPs in the deep WM of the lateral CB that contains the cerebellar nuclei (Cerrato et al., 2018). However, whether deep and lobule WM-NEPs are molecularly and functionally distinct is unknown. It is assumed, however that only NEPs in the WM of the lobules give rise to the INs that migrate to the molecular layer (ML) and populate it in an inside to outside manner (Figure 1A, B) (Brown et al., 2019; Sudarov et al., 2011). The full extent of the diversity of postnatal NEPs, and the molecular signature, lineage propensity and location of each NEP subtype are unknown.

**Figure 1.**
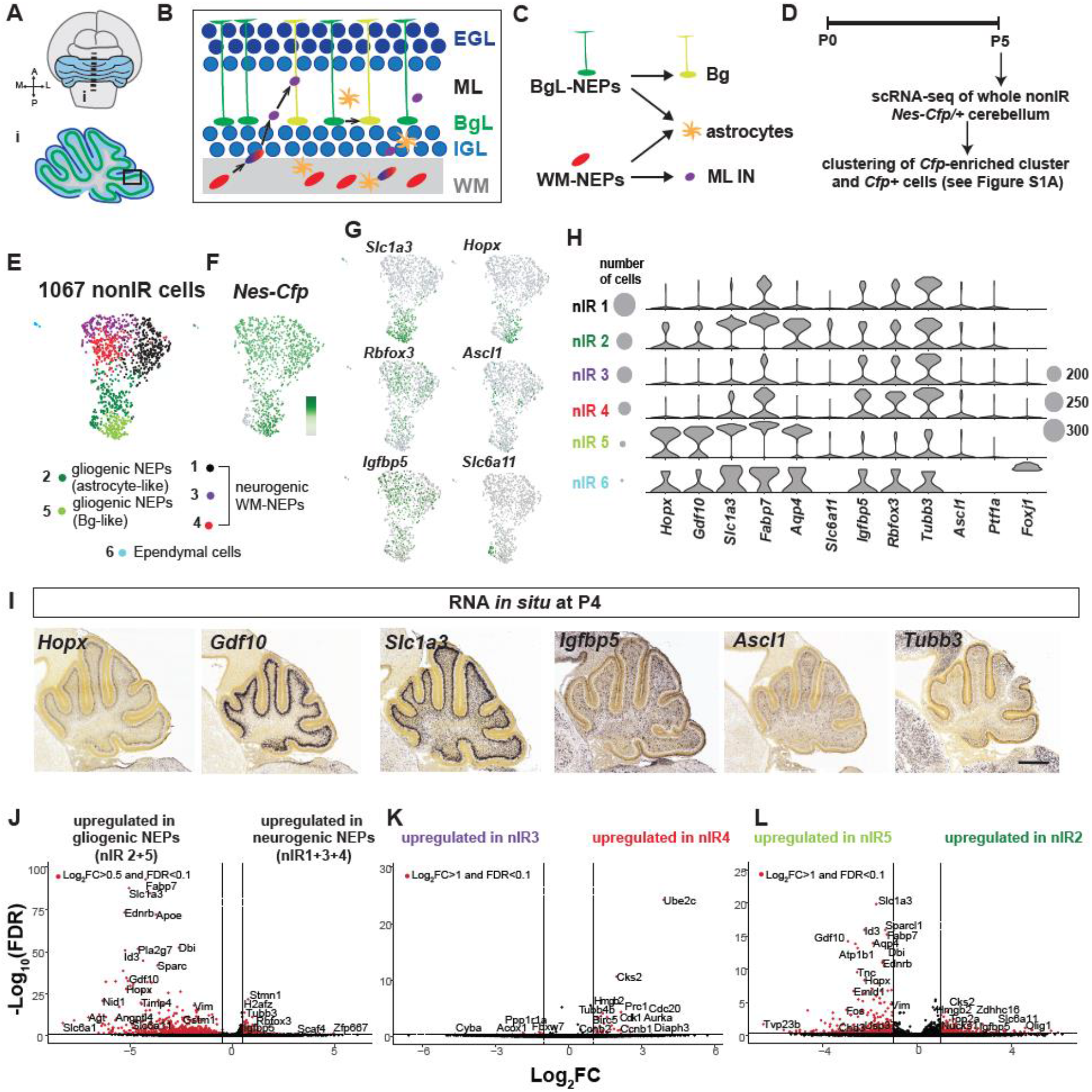
scRNA-seq of P5 cerebellar NEPs reveals diversity at steady state. **A-D.** Schematics showing the neonatal CB (A), cell types in each layer (B), the previously proposed lineages of the postnatal cerebellar progenitors (C) and the experimental plan (D). **E.** UMAP embedding of the clustering of 1,067 nonIR cells at stead state. **F-G.** Projections of the expression levels of *Nes-Cfp* and lineage markers on the UMAP (scale shows the log transformed CPM (counts per million) where the darker colors show higher expression). **H.** Violin plots showing the log_10_CPM expression levels of the canonical lineage genes and some of the top significantly enriched genes in clusters shown in E (Table S1). Scaled circles show the numbers of cells within each cluster. **I.** Allen Brain Atlas P4 RNA *in situ* hybridization data showing the expression patterns of marker genes identified by scRNA-seq. **J-L**. Volcano plots showing the differentially expressed genes (FC: fold change and FDR: false discovery rate) obtained from the pairwise comparisons of gliogenic- (nIR5+2) vs. neurogenic-NEPs (nIR1+3+4; J), nIR3 vs nIR4 (K) and nIR5 vs. nIR2 (L). Scale bars: 500μm

When the proliferating GCPs are depleted in the newborn CB either via genetic approaches or by irradiation, the EGL recovers within a week and the CB grows to almost a normal size (Altman and Anderson, 1971; Altman et al., 1969; Wojcinski et al., 2017; Wojcinski et al., 2018). A key aspect of the regenerative process involves BgL-NEPs, thought to be gliogenic, undergoing adaptive reprograming to replenish the lost GCPs. The first response of the BgL-NEPs is an increase in proliferation, followed by migration to the injured EGL where they initiate expression of *Atoh1* and become GCPs (Andreotti et al., 2018; Jaeger and Jessberger, 2017; Wojcinski et al., 2017; Wojcinski et al., 2018). Simultaneously, the WM-NEPs in the lobules reduce their proliferation and differentiation until the EGL is restored, leading to an overall delay in CB development, likely to ensure proper scaling of the different cell types of the cerebellar cortex (Wojcinski et al., 2017). Mosaic mutant analysis revealed that Sonic hedgehog signaling (SHH) is important for the adaptive reprograming of NEPs (Wojcinski et al., 2017). However, it is not known what molecular event induces the glial to neural fate change of BgL-NEPs upon injury and whether a new transitory cellular state facilitates the fate switch.

The commitment of neural stem cells to a neural fate is dependent upon a group of proneural basic helix-loop-helix (bHLH) transcription factors (Dennis et al., 2019; Guillemot and Hassan, 2017). In the CB, RL-derived excitatory neurons and VZ-derived inhibitory neurons are specified by distinct bHLH proteins typified by ATOH1 and ASCL1, respectively (RL-bHLH and VZ-bHLH). Fate mapping in the CB has shown that *Ascl1* is transiently expressed sequentially in a subset of VZ- and then NEP-derived cells that generate inhibitory neurons and is never involved in GCP production (Brown et al., 2019; Sudarov et al., 2011). Moreover, in cell culture ASCL1 is one of the primary proteins found to transform non-neural cells into neurons *in vitro*. For example, mis-expression of *Ascl1* along with *Brn2* and *Myt1l* converts fibroblasts into functional neurons, and ASCL1 acts as a pioneer factor that initiates the reprograming of a fate change via inducing a neurogenic transcriptional program (Vierbuchen et al., 2010; Wapinski et al., 2017). *In vivo*, ectopic expression of ASCL1 or other VZ-bHLH proteins is able to reprogram astrocytes to form neural cells upon injury (Grande et al., 2013; Heinrich et al., 2014; Liu et al., 2015; Zamboni et al., 2020). In contrast to the required mis-expression of VZ-bHLH proteins in all these studies, during GCP regeneration BgL-NEPs do not require ectopic expression of any genes to undergo a glial-to-neural fate switch upon injury, highlighting their high plasticity compared to other injury responsive glial cells outside the CB. However, whether and which proneural bHLH genes play a role in the acquisition of a neural fate by BgL-NEPs during GCP regeneration is unknown.

We used scRNA-seq in combination with genetic inducible fate mapping (GIFM) and loss of function studies to define molecularly distinct NEP populations in the postnatal developing CB and uncover a transcription factor involved in the fate switch of a VZ-derived gliogenic BgL-NEP to a RL-derived GCP. Five molecularly distinct NEP populations were identified at postnatal day (P) 5 and based on their lineage propensity they comprise 2 NEP subtypes. GIFM revealed that during homeostasis a *Hopx*-expressing NEP subtype is primarily gliogenic and resides in both the BgL and WM, whereas an *Ascl1*-expressing NEP subtype resides in the lobule WM and is neurogenic (produces INs). Furthermore, amongst the NEP subtypes seen after injury a new transitory state was identified in cells derived from *Hopx*-expressing BgL-NEPs and defined by *Ascl1* expression. Conditional mutagenesis showed that *Ascl1* is not only a marker of the transitory cell state but also plays a crucial role in GCP regeneration. Collectively, our results reveal the molecular diversity and the cellular plasticity of cerebellar NEPs and identify a context- (injury-) dependent transitory cellular state reponsible for a glial to neural fate switch that enables neonatal cerebellar regeneration.

## Results

### Identification of NEPs using scRNA-seq of whole P5 cerebellum during homeostasis and repair

In order to identify molecularly distinct NEP subtypes that are present in the postnatal CB and distinct cellular states that are acquired during adaptive reprogramming to become GCPs, we performed scRNA-seq on dissociated cells from 4 male P5 *Nes-Cfp*/+ pups that were irradiated (IR) at P1 or control littermates that were not irradiated (nonIR) (Figure S1A). An automated microwell-array-based platform was used to simultaneously profile 13,000 single cells from nonIR and IR animals (Yuan and Sims, 2016), followed by subclustering to identify pervasive and specific markers for molecularly distinct NEP subtypes (Figure S1A-C). Clustering of cells (6,000 nonIR, 7,000 IR, Figure S1D) using Phenograph (Levine et al., 2015) revealed the expected variety of cell types, with the greatest number of cells being neurons and the majority being GCPs and GCs based on the expression of *Atoh1* (GCP) and *Barhl1* (GCP/GC) (Figure S1D-G, Table S1). The other major cell types that were identified include astroglia (Bergmann glia, astrocytes, C7, Figure S1F), oligodendrocyte progenitors/oligodendrocytes, microglia and other parenchymal cells (Figure S1G). We used *Cfp* transcripts as a surrogate to identify cluster(s) containing NEPs and found *Cfp* expression was enriched only in the astroglial cluster C7 (Figure S1F). Interestingly, the *Cfp*+ cells in the astroglial cluster only represented 32% of all the *Cfp*+ cells, as scattered *Cfp*-expressing cells were observed in the neural clusters and the oligodendrocyte cluster (Figure S1G), suggesting that the transcript is retained in some of the immediate differentiated progeny of NEPs. Henceforth, we define NEPs as the cells in cluster 7 and all other *Cfp*+ cells present in the neural clusters (Figure S1A). Finally, our analysis of sections of P5 *Nes-Cfp*/+ cerebella showed that in addition to the BgL and the lobule WM, there are also CFP+ cells in the deep WM that houses the cerebellar nuclei (Figure S1H-J). These cells represent <10% of all the CFP+ cells in the CB, and therefore are likely poorly represented in our scRNA-seq data.

### scRNA-seq identifies two major subtypes and 5 populations of NEPs in the P5 CB at steady state

In order to determine whether distinct NEP subpopulations exist at steady state, we clustered the 1,067 NEPs from nonIR P5 mice (Figure 1D-F, Table S1). Phenograph analysis revealed five clusters (nIR1-5 for nonIR NEPs) and a small cluster of 13 ependymal cells (*Foxj1*+, nIR6) that cover the wall of the 3^rd^ ventricle (Figure 1G-H). Analysis of the Allen Brain Atlas RNA *in situ* mouse P4 data set for the significantly enriched genes in each cluster and lineage markers allowed further validation of cluster identities. nIR5 is enriched for Bg lineage genes (*Hopx, Gdf10*) (Carter et al., 2018; Heng et al., 2017; Mecklenburg et al., 2014), nIR2 also is enriched for these genes but has lower expression of them and both nIR2 and nIR5 express pan-astrocyte genes (*Slc1a3, Aqp4, Fabp7*) (Figure 1I). One gene enriched specifically in nIR2 is *Slc6a11,* an astrocyte specific GABA transporter (Boddum et al., 2016). Interestingly, *Slc6a11* was expressed in the subset of nIR2 cells that do not express *Cfp*, indicating that they are differentiated astrocytes and the rest of the cluster might be astrocyte progenitors (Figure 1G,H). On the other hand, the high expression of Bg genes in nIR5 could indicate the cells are Bg progenitors. Three clusters (nIR1, 3, 4) showed similar patterns of expression of neural genes (e.g. *Rbfox3, Tubb3*) and a gene enriched in the WM (*Igbfbp5*) based on RNA *in situ* analysis (Allen Brain Atlas, Figure 1G-I). Interestingly, the proneural VZ-bHLH gene, *Ascl1,* was enriched in nI4, although it is present in all three clusters at low levels (Figure 1G-I). Other immature IN marker genes such as *Pax2* were not detected at high levels in our dataset. Furthermore, differential expression analysis of the genes expressed in the gliogenic clusters (nIR2, 5) compared to neurogenic clusters (nIR1, 3, 4) highlight further the gliogenic and neurogenic gene signature of each NEP subtype (Figure 1J). In summary, scRNA-seq analysis identified two major NEP subtypes, gliogenic (nIR2, 5) and neurogenic (nIR1, 3, 4), which appear to be further subdivided but into less distinct populations.

To identify differences between the neurogenic clusters (nIR1, 3 and 4) we performed differential expression analysis for pairwise comparisons of the three clusters. The results further highlighted the similarities between nIR1, 3 and 4, as there were few significantly differentially expressed genes (FDR<0.1, Log_2_FC>1, n=39 genes nIR1 vs. nIR3, n=16 genes nIR1 vs. nIR4 and n=28 genes nIR3 vs. nIR4, Figure 1K) and the genes were related to cell cycle (Table S2, Figure 1K). The results indicate that nIR1 represents a more proliferative population, consistent with its slightly higher *Cfp* expression, whereas the nIR4 and nIR3 could represent NEP populations transitioning into a more differentiated state.

Differential expression analysis was also performed on the two gliogenic clusters and confirmed that nIR5 has significantly higher expression of Bg associated genes (*Hopx* and *Gdf10*), supporting that nIR5 represents Bg progenitors and their immediate Bg progeny, whereas nIR2 expresses astrocyte genes at a higher level and thus could represent astrocyte progenitors and mature astrocytes (Figure 1L). nIR2 also shows expression of *Rbfox3* (and *Tubb3*) in some cells and the WM enriched *Igfbp5* gene (Figure 1G,H). Therefore nIR2 could include a more plastic group of NEPs including ones in the WM that can form astrocytes and possibly INs (Figure 1G,H). Collectively, our data show that the primary molecular signatures that distinguish NEP subpopulations is based on their lineage propensities, gliogenic vs neurogenic subtypes. In addition, neurogenic NEPs exhibit additional molecular diversity seemingly associated with their differentiation state, whereas the two gliogenic NEPs populations could be dedicated to producing Bg (nIR5) or astrocytes (nIR2).

### Ascl1- and Hopx-expressing NEPs have distinct cell lineages

In order to confirm the identities of the NEP subtypes identified via scRNA-seq and study their lineages, we performed GIFM for the neurogenic-NEPs using an *Ascl1*^*CreERT2*^ allele and for the gliogenic-NEPs using a *Hopx*^*CreERT2*^ allele in combination with a *R26*^*lox-STOP-loxTdTomato*^ reporter (*Ascl1-TdT* and *Hopx-TdT*, mice respectively). When Tamoxifen (Tm) was administered at P5 to label cells at a similar time point as our scRNA-seq data (Figure 2A), short-term GIFM (analysis 2 days after Tm injection) in *Ascl1-TdT* mice showed that the majority of labeled cells were restricted to the lobule WM at P7 with scattered cells in the IGL that had the appearance of migrating immature ML INs and interestingly no cells in the deep WM (Figure 2B-E, Figure 2SA-D). Double labeling with the immature IN marker PAX2 confirmed that most of the TdT+ cells in the WM are INs (Figure S2I). Analysis of long-term GIFM (P30) showed that almost all of the progeny of the *Ascl1*+ WM-NEPs are ML INs (97.9 ± 2.3 %, n=5), specifically the stellate cells which are the latest born INs located in the outer ML (Figure 2F-I). Only rare astrocytes in the lobule WM or IGL and no GCs were labeled in the adult CB. These results not only confirm there is a neurogenic NEP subtype in the lobule WM, but that it has a restricted lineage and at P5 gives rise almost exclusively to Stellate INs.

**Figure 2.**
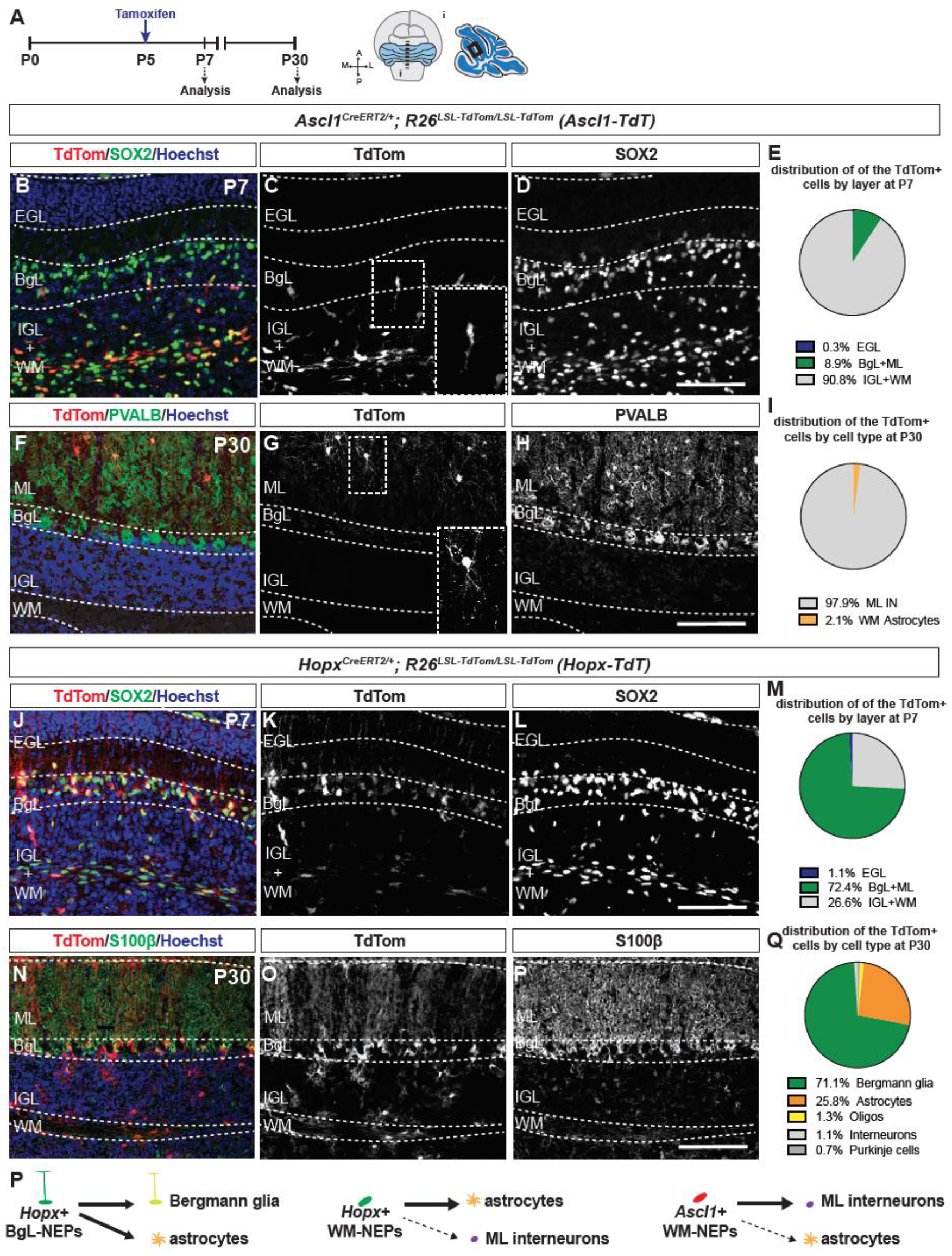
GIFM identifies the lineages of the *Ascl1*+ neurogenic-NEPs and the *Hopx*+ gliogenic-NEPs. **A.** Experimental plan and the region of the CB shown in the images. **B-I.** IF analysis used for the quantification of the distributions of the TdT+ cells by layer at P7 (B-E) or by cell type at P30 (F-I) in *Ascl1-TdT* animals (n=3 brains/age). Insets show the morphology of a migrating IN (C) progenitors and a stellate neuron (G). **J-M.** IF used for analysis and the quantification of the distributions of the TdT+ cells by layer at P7 (J-M) or by cell type at P30 (N-Q) in *Hopx-TdT* animals (n=3 brains/age). **P.** Proposed summary of the linages generated from the *Ascl1-* and *Hopx-*expressing NEPs at P5. Scale bars: 50μm

Short-term labeling in P7 *Hopx-TdT* animals showed that all of the TdT+ cells were in the lobules and the majority were in the BgL (72.4 ± 6.2%, n=3, Figure 2J-Q and S2E-H). Marker analysis showed that both NEPs (SOX2+/S100β−) and Bg (SOX2+/S100β+) in the BgL were TdT+ (Figure S2J). Curiously, a subpopulation of lobule WM-NEPs was also TdT+ (26.6 ± 6.1%, n=3) along with very rare SOX2+ cells in the inner EGL (1.1 ± 0.3%, n=3, Figure 2J-M). Analysis of the *Hopx-TdT* mice at P30 showed that 98.2 ± 0.6% (n=3) of the TdT+ cells were astroglia (71.1 ± 2.7% Bg, 25.8 ± 2.1% astrocytes in the IGL and WM and 1.3 ± 1.0 % oligodendrocytes in the WM) and only ~1.8% were INs (1.10 ± 0.3 % in the ML) (Figure 2N-Q) and no labelled cells were seen in the deep WM (Figure SE-H).

RNA *in situ* analysis of *Igfbp5* showed expression in the deep and lobule WM, whereas *Hopx* and *Ascl1* expression were restricted to the lobules at P5 (Figure S2M-O). As there was no labeling in the deep WM cells at both P7 and P30 both in the *Ascl1-TdT* and the *Hopx-TdT* cerebella, this reveals that the lineages of NEPs expressing these genes are restricted to the lobules (Figure 2SA-H). Given that the INs of the cerebellar nuclei are generated during embryonic development directly from the VZ, the deep WM-NEPs should only be gliogenic but a specific marker gene is yet to be determined. Overall the GIFM studies validate the identities of the gliogenic- vs. neurogenic-NEP subtypes obtained via scRNA-seq (Figure 1E), uncover a WM-NEP population in the lobules that expresses *Hopx* and demonstrate that *Ascl1*-expressing NEPs in the neurogenic clusters have an ML IN-restricted lineage at P5 (Figure 2P).

### Irradiation induces changes in the NEP populations and the emergence of an Ascl1+ transitory NEP

The P5 time point was selected for scRNA-seq because it should allow a variety of cellular states during regeneration to be identified since in IR mice the NEPs and their immediate progeny that express CFP are undergoing all stages of adaptive reprogramming, including proliferation, a fate change, and migration to the EGL (Figure S3A, Figure 3A-D). When the 1,472 NEPs from the P5 IR cerebella were analyzed using Phenograph, 6 clusters were obtained (IR1-6, for IR NEP clusters, Figure 3E, Figure S1A,C, Figure S3B-G, Table S1). Gene set enrichment analysis (GSEA) of the differentially expressed genes between nonIR and IR NEPs, ascertained by single cell differential expression analysis (SCDE) (Kharchenko et al., 2014), identified many cellular processes altered 4 days after injury, including upregulation of neural fate commitment and glial differentiation in the IR NEPs compared to increased synapse maturation in the nonIR NEPs (NES>1.5, p<0.05, Figure S3H, Table S3).

**Figure 3.**
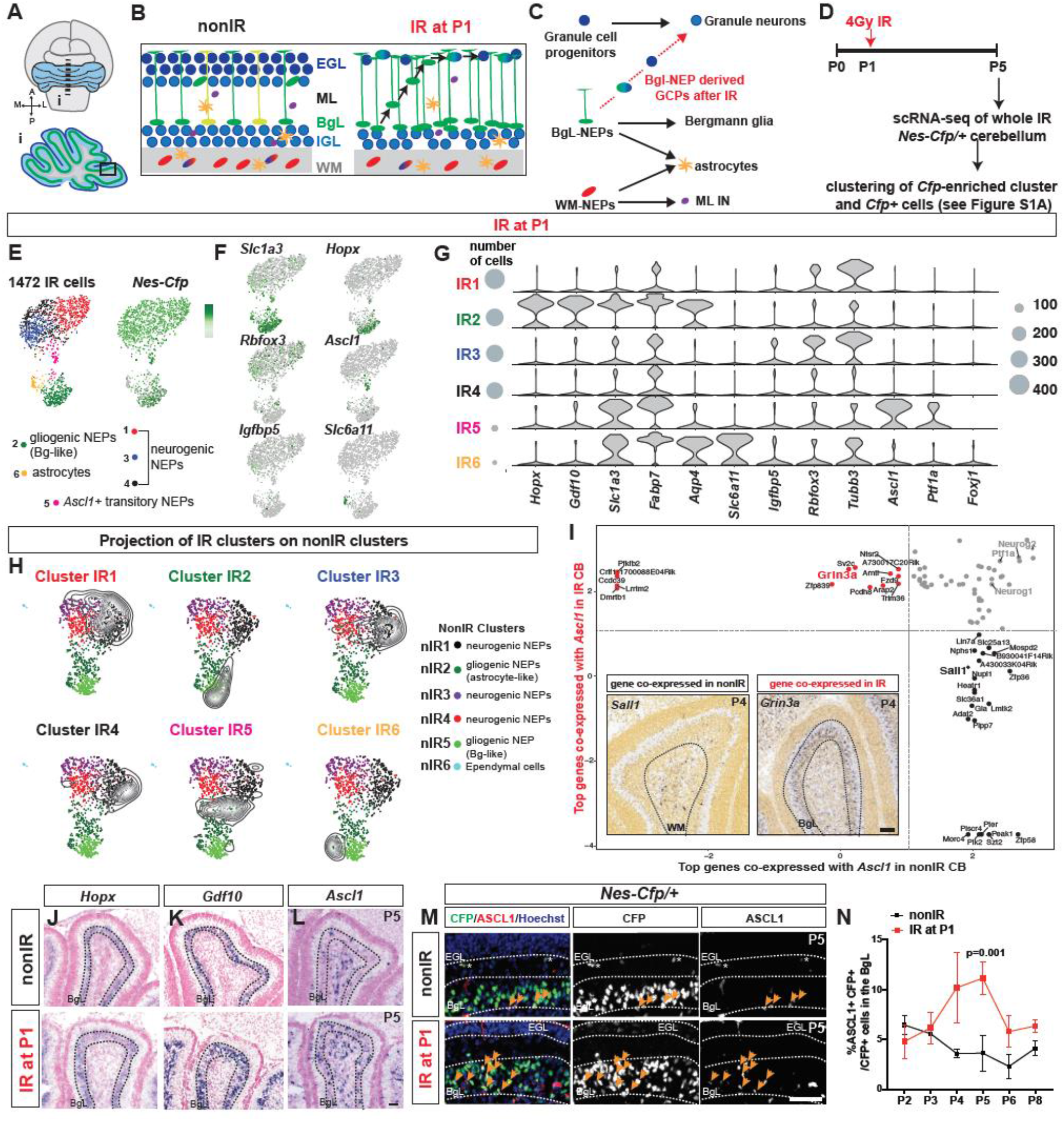
scRNA-seq reveals changes in NEP populations and emergence of an *Ascl1*+ transitory cell state upon injury. **A-D.** Schematics showing the CB (A), cell types in each layer of the P5 neonatal CB and the changes that occur after IR at P1 (B), the previously proposed changes in the lineages of the postnatal cerebellar progenitors (C) and the experimental plan (D). **E.** UMAP embedding of the clustering of 1,476 IR NEPs. **E-F.** Projections of the expression levels (log transformed CPM (counts per million) where the darker colors show higher expression) of the *Nes-Cfp* transcripts and other lineage markers on the UMAPs. **G.** Violin plots showing the log_10_CPM expression levels of the canonical lineage genes and some of the top significantly enriched genes in nonIR (nIR) clusters (Table S1). Scaled circles show the numbers of cells. **H.** Projections of IR NEP clusters (density plots, the black lines indicate low and grey lines indicate high density) on to the nonIR UMAP embeddings (dot plots, Figure 1E). **I.** Analysis of the genes that are most likely to be co-expressed with *Ascl1* in the nonIR and IR NEPs. Insets show Allen Brain Atlas RNA *in situ* analysis for *Sall1* and *Grin3a* in the P4 CB. **J-L.** RNA *in situ* analysis of P5 nonIR and IR brains for BgL-NEP (*Hopx, Gdf10*) and neurogenic-NEP (*Ascl1*) markers. **M**. IF analysis for ASCL1 and CFP on P5 nonIR and IR *Nes-Cfp* cerebella. **N.** Quantification of percentage of ASCL1+ CFP+ cells in the BgL at P2-6 and P8 in nonIR and IR *Nes-Cfp* cerebella (Two-way ANOVA, F_(1,15)_=25.75, p=0.0001). Scale bars: 50μm

The clustering of the IR NEPs, as with the nonIR NEPs, was dictated by a gliogenic or neurogenic transcriptional signature. In addition, 2 gliogenic clusters (IR2 and 6, Figure 3E-G) and 3 neurogenic clusters (IR1, 3 and 4, Figure 3E-G) were again identified. However, importantly an additional distinct cluster was identified in IR NEPs (IR5). Most interestingly, the new cluster in the IR NEPs (IR5) showed enrichment for the neurogenic VZ-bHLH factor *Ascl1* as well as glial genes such as *Slc1a3* and *Fabp7* (Figure 3F-G). In terms of the gliogenic clusters, IR2 showed enrichment for Bg genes (*Hopx* and *Gdf10*) similar to nlR5, whereas IR6 cells were enriched for astrocyte genes (*Slc1a3*, *Aqp4* and *Fabp7*) but not Bg genes and were mainly *Cfp*- (Figure 3E-G). Although such a distinct cluster was not detected in nonIR NEPs, a group of similar mature astrocytes was identified as a subgroup within nIR2 (Figure 1E,H). RNA *in situ* analysis of the BgL-NEP markers *Gdf10* and *Hopx* on sections from nonIR and IR P5 cerebella showed that their expression pattern was not changed by IR, as expression remained in the BgL, although as expected the BgL was thicker reflecting the increased proliferation of BgL-NEPs upon irradiation (Figure 3J-K) (Wojcinski et al., 2017). In terms of the neurogenic clusters (IR1,3 and 4), the WM-NEP marker *Igfbp5* was expressed in a smaller proportion of the IR NEPs (270/1067 nonIR NEPs, Figure 1G vs. 137/1472 IR NEPs, Figure 3F). RNA *in situ* analysis of sections from IR P5 cerebella validated the reduction in *Igfbp5* expression in both the lobule and deep WM cells after IR compared to nonIR controls (Figure S3I-M).

To further investigate how the IR NEP clusters are related to the nonIR NEP subtypes, we projected the IR cells (density plots) onto the nonIR (dot plots) UMAP embedding (Figure 3H). This analysis led to multiple interesting observations. First, the 3 IR neurogenic-NEP clusters (IR1, 3 and 4) overlapped specifically with nIR1, which appears to represent the most proliferative and least differentiated (*Cfp* high) steady-state neurogenic-NEP subpopulation (Figure 3H). Secondly, IR2 overlapped with nIR5, the gliogenic NEP cluster with Bg characteristics, suggesting that these NEPs primarily generate Bg. Consistent with the astrocyte expression signature of IR6, it overlapped specifically with the subregion of nIR2 that has *Cfp-* astrocytes (Figure 3H). It is curious that very few IR cells in IR2 or IR6 projected to the subregion of nIR2 that does not contain astrocytes and contains progenitors. Moreover, thirdly, there was no clear nonIR counterpart to lR5, which projected between nIR2 and the three neurogenic-NEPs clusters (Figure 3H). Thus, IR5 could represent a transitory cell population induced upon injury with a new gene expression signature that shares some similarities with the nonIR Bg-like gliogenic-NEPs (nIR2) and the neurogenic-NEP subtype. Differential expression analysis of IR5 compared to gliogenic- (IR2) or neurogenic- (IR1, 3 and 4) NEPs further revealed IR5 has aspects of both the VZ-bHLH neurogenic and glial signatures (Figure S3N-Q, Table S4). Finally, as a negative control, no ependymal cells were detected in the IR NEP dataset (based on *Foxj1* expression), and none of the IR NEPs projected onto the small cluster of ependymal cells observed in the nonIR data (nIR6).

### Transcriptional programs of *Ascl1*+ NEPs diverge between nonIR and IR conditions

We next investigated the transcriptional differences between all the *Ascl1*+ cells in nonIR compared to IR NEPs. We assessed co-expression and mutual exclusivity for all gene pairs and calculated probability ratios of detecting two genes in the same cell (Mizrak et al., 2019) for the top 50 genes with high *Ascl1* co-expression probability ratios (15/100 showed high co-expression probability ratios with *Ascl1* in both IR and nonIR cells, Table S5). Among the genes co-expressed with *Ascl1* in both conditions were three other proneural VZ-bHLH genes, *Ptf1a* and *Neurog1/2,* (Figure 3I). Interestingly, we also detected sets of genes that were co-expressed with *Ascl1* only in nonIR or in IR NEPs (Figure 3I). Analysis of P4 RNA *in situ* data (Allen Brain Atlas) for *Sall1* that is co-expressed with *Ascl1* in nonIR-NEPs revealed its expression is restricted to the WM where *Ascl1*+ NEPs normally reside (Figure 3I). Interestingly, *Grin3a* that is co-expressed with *Ascl1* in IR NEPs is expressed in the BgL, consistent with the idea that after irradiation *Ascl1* is induced in some BgL-NEPs (Figure 3I). Furthermore, some of the other genes co-expressed with *Ascl1* in the IR NEPs, *Grin3a*, *Arntl* and *S1pr3* have been implicated in astrocyte injury-related functions (Lananna et al., 2018; Rothhammer et al., 2017; Rusnakova et al., 2013), indicating the new cluster derives from a gliogenic NEP subtypes. These results further highlight that the *Ascl1*+ NEPs in the IR cerebella have a glial gene signature, leading us to suggest that the injury responsive gliogenic BgL-NEPs turn on *Ascl1* upon injury.

To test whether injury does indeed induce VZ-bHLH gene expression in BgL-NEPs, we performed RNA *in situ* hybridization for *Ascl1* on P5 nonIR and IR cerebellar sections and observed a clear increase in expression of *Ascl1* in the BgL (Figure 3L). IF analysis of P5 *Nes-Cfp*/+ cerebella also confirmed an increase in the number of ASCL1+ CFP+ cells in the BgL (3.1 ± 0.4 –fold, n=3) of IR cerebella compared to nonIR P5 littermates (Figure 3M). Interestingly, ASCL1 expression was not observed in most of the CFP+ cells in the EGL indicating that *Ascl1* is transiently induced upon injury in BgL NEPs prior to migration to the EGL (Figure 3M). Finally, quantification of the number of ASCL1+ cells every 24 hours between P2 and P6 and at P8 showed that expression in BgL-NEPs (CFP+ cells) peaks 3-4 days after IR (Figure 3N). Based on our scRNA-seq comparison of P5 IR and nonIR NEPs and *in vivo* identification of the location of the NEP populations, we propose that that upon injury, normally gliogenic BgL-NEPs turn on a new transcriptional program involving proneural VZ-bHLH transcription factors that allows a switch to a neural fate to occur prior to their mobilization to the EGL.

### ASCL1+ transitory cells and granule cells are derived from Hopx-NEPs after injury

In order to demonstrate that *Hopx-*derived BgL-NEPs give rise to ASCL1+ cells that produce GCPs and GCs after irradiation, we performed GIFM with *Hopx-TdT* animals. When Tm was administered at P0 (Figure 4A), analysis of P7 cerebella revealed a significant increase in the percentage of ASCL1+ TdT+ cells of all TdT+ cells in the BgL in IR pups (12.7 ± 3.1%, n=3) compared to nonIR littermates (1.4 ± 1.4%, n=3, Figure 4B-H). In the nonIR brains, as expected ASCL1 expression was restricted to the WM and IGL and showed almost no overlap with *Hopx*-derived TdT+ cells in the WM (asterisks, Figure 4B-G). Interestingly, in IR pups the percentage of the ASCL1+ TdT+ double positive cells amongst all the ASCL1+ cells in the BgL was higher than the percentage of SOX2+ TdT+ double positive cells amongst all the SOX2+ cells (a pan-NEP/Bg/astrocyte marker) in the BgL (~39.2±7.7% in IR and 6.5 ± 1.9% in nonIR vs 15.5 ± 6.7% IR and 26.1 ± 8.8% in nonIR, respectively, n=3, Figure 4I). This result suggests that *Hopx*^*CreERT2*^ preferentially marks the BgL-NEPs at ~P1 with the ability to undergo adaptive reprograming in response to loss of GCPs. Importantly, TdT+ cells were present in the EGL of the IR P7 cerebella, and most expressed SOX2, indicating that progeny of SOX2+ *Hopx-TdT*+ labeled BgL-NEPs migrated to the site of injury (Figure 4J-P). At P30, we observed a significant increase in the density of TdT+ cells in IR *Hopx-TdT* brains compared to nonIR (Figure 4Q-S), which was primarily driven by a significant increase in the density of TdT+ GCs in the IGL (Figure 4S-T). Although when Tm was given at P0 compared to P5 in *Hopx-TdT* mice we found a higher proportion of the TdT+ cells to be WM-NEPs at P7 (49.1% vs. 26.6%, Tm at P0 vs. P5, n=3) and ML INs at P30 (31.4% vs. 1.1%, Tm at P0 vs. P5, n=3), there was no significant difference in the density of TdT+ ML INs between IR and nonIR mice (Figure 4S-T). Finally, no TdT+ cells were detected in the deep WM of IR or nonIR *Hopx-TdT* animals that were given Tm at P0 (Figure S4A-B). Overall, these results provide experimental evidence that *Hopx*+ BgL-NEPs give rise to ASCL1-expressing transitory cells that change their fate and become GCPs and then GCs.

**Figure 4.**
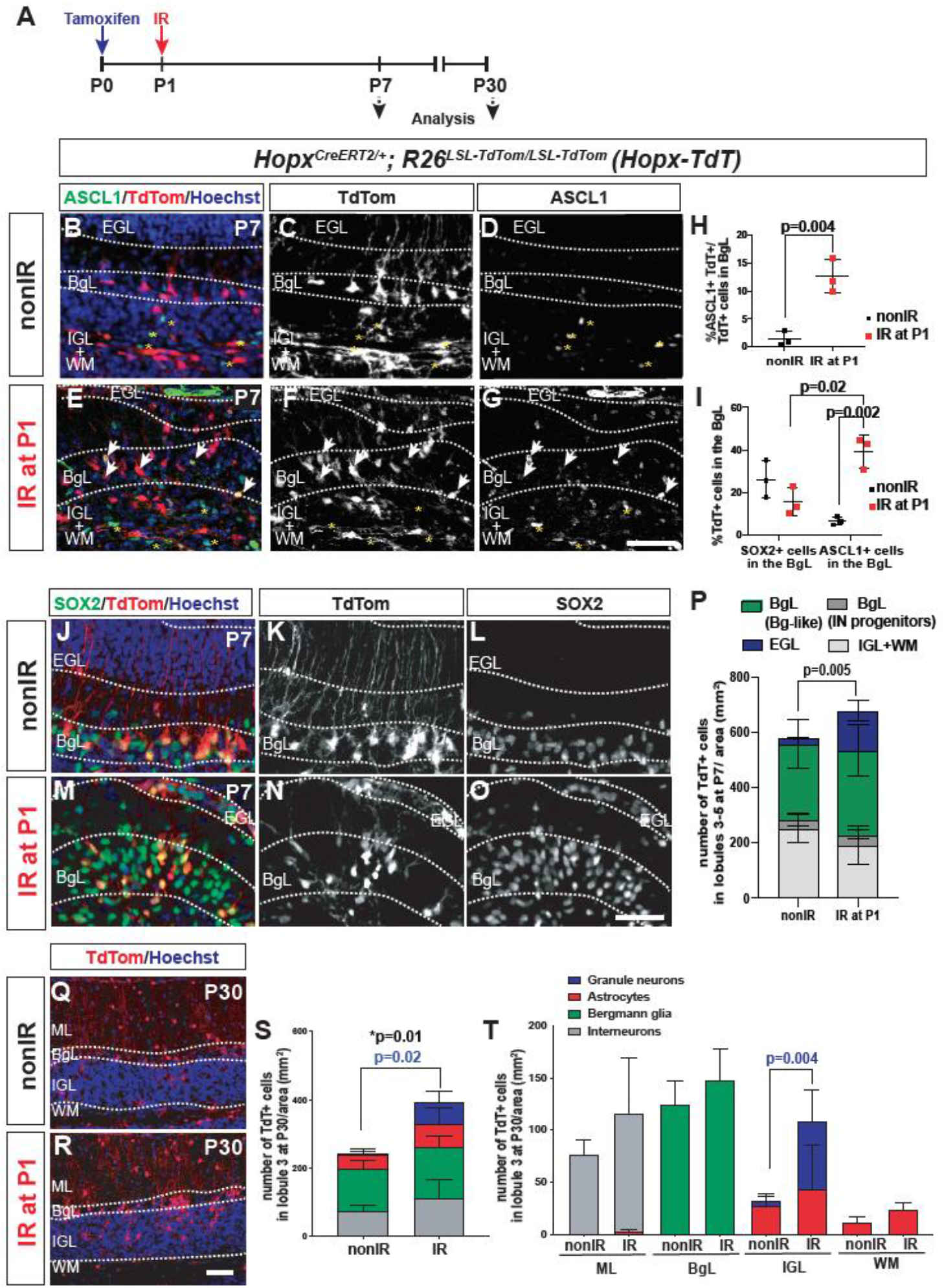
*Hopx*-derived BgL-NEPs give rise to GCPs via an ASCL1+ transitory cell state. **A.** Experimental plan. **B-H.** IF analysis (B-G) and quantification (H) of the percentage of the TdT+ and ASCL1+ cells per TdT+ cells in the BgL at P7 on sections from *Hopx-TdT* nonIR (B-D) and IR (E-H) brains (n=3 brains, Student’s t-test, p=0.004). **I.** Quantification of the percentage of *Hopx*-derived TdT+ cells in all BgL-NEPs and Bg (SOX2+ cells) or in all *Ascl1*+ BgL-NEPs (n=3/condition, Two-way ANOVA, F_(1,8)_=7.928, p=0.02). **J-O.** IF analysis of P7 nonIR (J-L) and IR (M-O) *Hopx-TdT* brains. **P**. Quantification of the density of TdT+ cells in nonIR and IR *Hopx-TdT* brains at P7 (n=3/condition, EGL: Student’s t-test, p=0.005). **Q-R** IF analysis of P30 nonIR (J-L) and IR (M-O) *Hopx-TdT* brains. **S-T.** Quantification of the density of the progeny of nonIR and IR *Hopx-TdT* brains at P30 by cell types (S, n=3/condition, granule cells: Student’s t-test, p=0.02) and by the layer of the CB (T) (n=3/condition, Two-way ANOVA, F_(7,42)_=8.718, p<0.0001). *p-value in (R) represents the change in the total number of cells (n=3/condition, Student’s t-test). Scale bars: 50μm

### ASCL1 marks a neurogenic BgL-NEP population during adaptive reprogramming that gives rise to GCs

We next tested whether *Ascl1* marks a new BgL-NEP transitory cellular state after irradiation that generates GCs by analyzing nonIR and IR *Ascl1-TdT* animals given Tm at P5 (Figure 5A), the stage with the largest number of injury-induced ASCL1+ BgL-NEPs (Figure 3N). Analysis of P6 and P7 cerebella revealed a population of TdT+ BgL-NEPs that are SOX2+/S100β− and have glial fibers projecting to the pial surface only in pups irradiated at P1 (Figure 5C,E,G,I). In contrast, the few TdT+ cells in the BgL of nonIR cerebella had a morphology of migrating progenitors (Figure 5B,D,F,H). Quantification of the density of TdT+ cells at P7 revealed a significant increase in the density of TdT+ cells in the EGL and BgL (Figure 5J). Furthermore, the number of labeled cells in the EGL increased significantly between 24 and 48 after Tm in IR cerebella (Figure 5C, J). Analysis of the proliferation of the TdT+ cells showed that in the IR pups, *Ascl1-TdT*+ cells in the BgL (both BgL-like and IN progenitor-like) and EGL incorporated EdU, as well as the *Ascl1-TdT*+ cells in the IGL+WM (Figure S4C-D). On the other hand in the nonIR brains, *Ascl1-TdT*+ IN progenitors in the IGL+WM and some that started migrating outwards towards the ML incorporated EdU, suggesting that all the *Ascl1*-expressing nonIR and IR NEPs are proliferative (Figure S4C-D).

**Figure 5.**
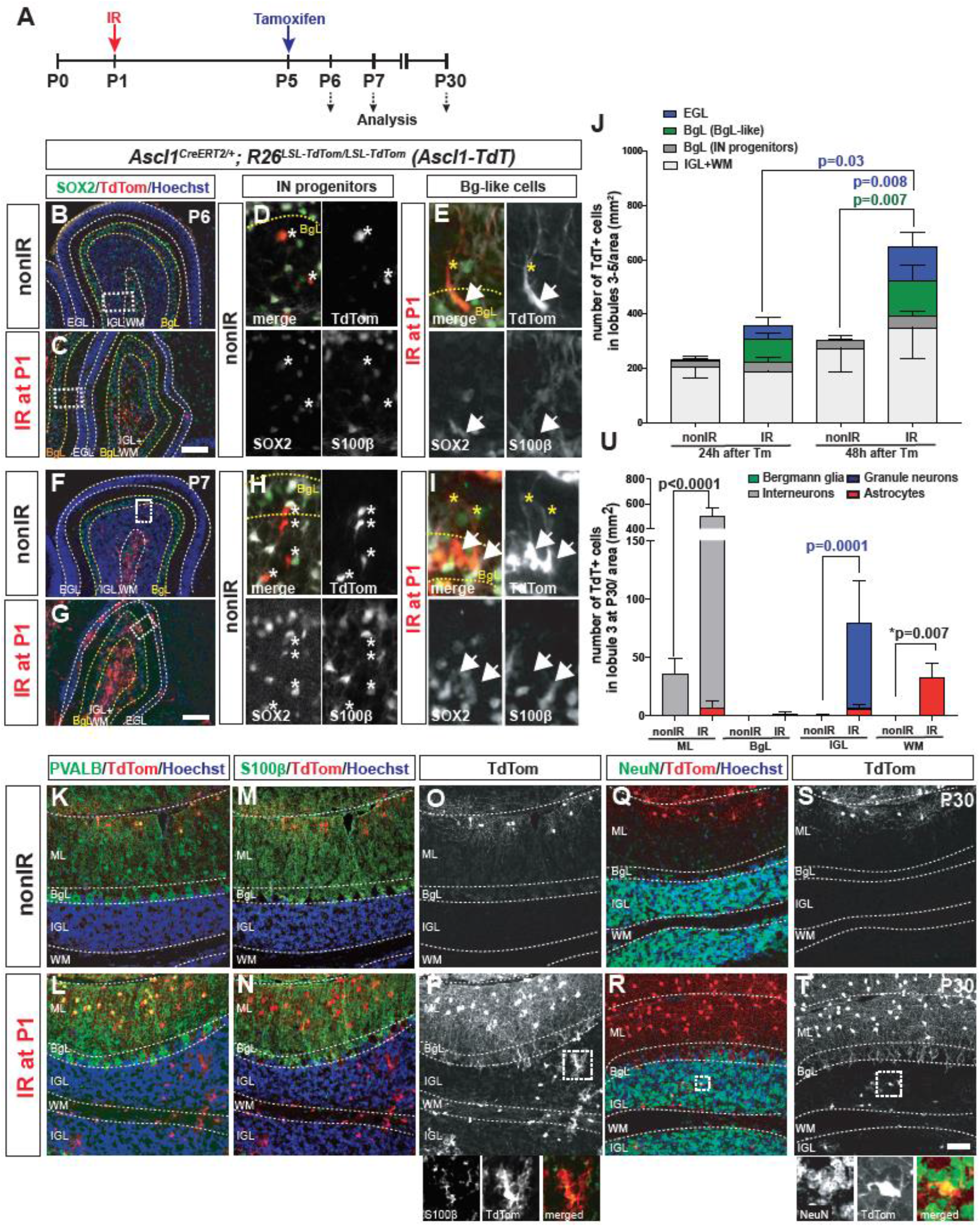
Injury induced *Ascl1*+ BgL-NEPs give rise to GCs and astrocytes but not Bg. **A.** Experimental plan. **B-I.** IF analysis of *Ascl1-TdT* brains at P6 (24h after Tm at P5, B-E) and at P7 (48h after Tm, F-I). Insets show a migrating IN progenitors (asterisks; D, H) and a SOX2+ S100β− BgL-like *Ascl1-TdT* cell (arrows; E, I). Yellow asterisks show the pial projection of the TdT+ cells. **J.** Quantification of the density of the TdT+ cells in the different layers of the CB at P6 and P7 in nonIR and IR *Ascl1-TdT*+ mice given Tm at P5 (BgL-like cells were identified by their pial projections (n=4, Two-way ANOVA, all layers: F_(3,32)_=12.89, p<0.0001, EGL: F_(1,8)_=23.75, p=0.001) **K-T**. IF analysis of nonIR and IR P30 *Ascl1-TdT* ceerebella showing examples of the TdT+ ML INs (PVALB), astrocytes (S100β) and GCs (NeuN in the IGL). **U.** Quantification of the density of the TdT+ cells by cell type in the different layers of the CB of P30 nonIR and IR *Ascl1-TdT* animals (n=3/condition, Two-way ANOVA, F_(7,14)_=137.9, p<0.0001, WM astrocytes: Student’s t-test, p<0.007). Scale bars: B-G: 100μm, K-T: 50μm

Analysis of P30 IR *Ascl1-TdT* cerebella revealed several key findings. Firstly, the P30 IR brains had a significant and large increase in the density and the proportion of the TdT+ cells that were GCs (13.8 ± 2.0 % of all TdT+ cells, n=4 IR compared to 1.0 ± 1.4% of TdT+ cells, n=5 nonIR, Figure 5R,T,U). Secondly, almost no TdT+ Bg were detected in the P30 IR *Ascl1-TdT* cerebella, suggesting that *Ascl1* induces the BgL-NEPs to switch from making Bg to GCPs and that the switch is irreversible (Figure 5M-N, U). Thirdly, there was a delay in production of ML INs after irradiation, as demonstrated by an increase in the density of TdT+ ML INs per lobule that included both earlier born Basket INs and later born Stellate INs (Figure 5K-L, U, Figure S4E-G). Finally, we observed an increase in astrocyte production in the IR *Ascl1-TdT* brains, primarily in the WM (Figure 5M-P, U). Surprisingly, after irradiation we detected an emergence of TdT+ astrocytes in the deep WM in *Ascl1-TdT* cerebella at P30, unlike in nonIR *Ascl1-TdT* cerebella or IR and nonIR *Hopx-TdT* brains where no TdT+ deep WM astrocytes were detected (Figure S4A-B, H). Since no TdT+ cells were detected in the deep WM of IR *Ascl1-TdT* cerebella at P6 and P7, this indicates that the TdT+ cells must originate from the lobules (Figure S3H). This result shows that two different NEP subtypes (*Ascl1*+ vs *Hopx*+) have different lineage responses to injury.

Finally, live imaging of lobule 3 in thick cerebellar sections from P7 cerebella showed that the *Ascl1-TdT*+ cells do indeed migrate from the BgL to the EGL in the IR cerebella, whereas in nonIR P7 slices no cells migrate from BgL to EGL (Figure S4A-C, and movie 1-2). Collectively, these results demonstrate that GCPs are generated from an *Ascl1*-expressing Bgl-NEP upon injury. Furthermore, irradiation leads to a delay in production of INs and ectopic astrocyte production, likely from normally neurogenic WM-NEPs.

### Ascl1 is required in Hopx+ NEPs for adaptive reprogramming and production of GCs

Given the surprising upregulation of *Ascl1* in the normally gliogenic BgL-NEPs upon death of GCPs, we tested whether *Ascl1* is required for repair of the GC lineage after cerebellar irradiation using a conditional knockout (CKO) approach. *Hopx*^*CreERT2*/+^; *Ascl1*^*fl/fl*^ animals injected with Tm at P0 (*Hopx*-*Ascl1* CKOs) were used to delete *Ascl1* primarily from BgL-NEPs and their regenerative capacity was compared to *Ascl1*^*fl/fl*^ littermate controls after irradiation at P1 (Figure 6A). Analysis of P30 cerebella showed a significantly greater reduction in cerebellar area at the midline in IR *Hopx*-*Ascl1* CKOs compared IR controls (Figure 6B-E, J). Furthermore, the IGL was less organized in some lobules of the IR mutants than in controls, further demonstrating impaired regeneration (Figure 6F-I). In order to rule out that the impairment of the regeneration is due to a loss of one copy of the *Hopx* gene in the *Hopx*-*Ascl1* CKO animals, we compared cerebellar areas of IR and nonIR *Hopx-TdT* animals and their *R26*^*TdT/*+^ littermates and found no significant changes between the genotypes, showing that loss of one copy of *Hopx* does not impair regeneration (Figure S6A-D, I). Interestingly, the loss of one copy of *Ascl1* resulted in a mild but significant reduction in the area of the CB of *Ascl1-TdT* animals compared to *R26*^*TdT*/+^ littermate controls, further confirming that ASCL1+ is required for regeneration after irradiation (Figure S6E-H, J).

**Figure 6.**
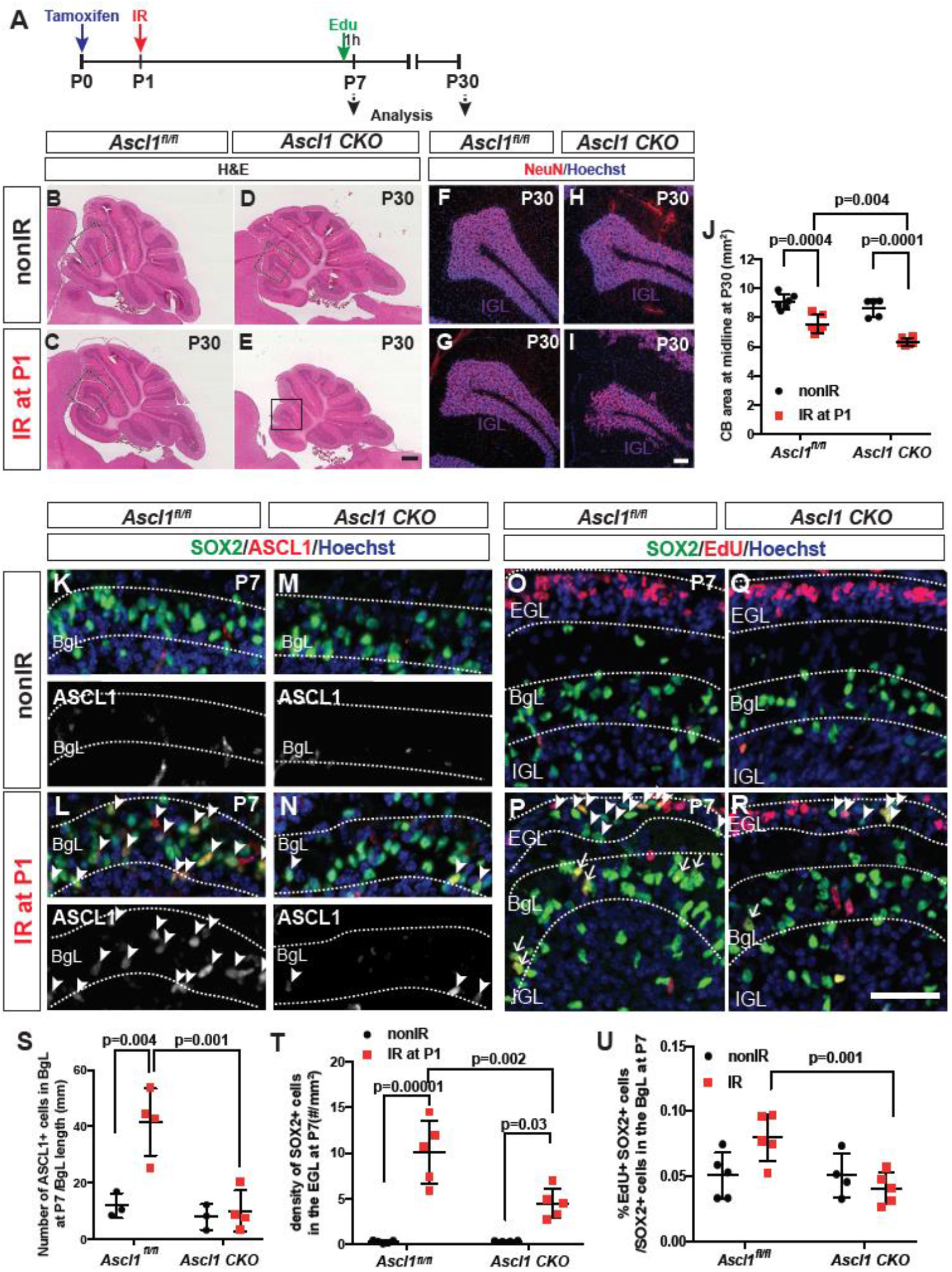
*Ascl1* is required for the adaptive reprograming of BgL-NEPs into granule cells. **A.** Experimental plan. **B-E.** H&E images of midsagittal sections from P30 nonIR and IR control or *Ascl1* CKO cerebella. **F-I.** IF analysis of NeuN at P30. **J.** Quantification of midline cerebellar area (n≥5/condition, Two-way ANOVA, F_(1,19)_=14.39, p<0.001). **K-N.** IF analysis for ASCL1 on sections from P7 nonIR and IR control and *Ascl1* CKO cerebella (arrowheads: ASCL1+ BgL-NEPs). **O-R**. IF analysis of SOX2 and EdU on sections from P7 nonIR and IR control and *Ascl1* CKO cerebella. **S-U.** Quantification of the density of ASCL1+ cells in the BgL (S, n≥3/condition, Two-way ANOVA, F_(1,10)_=16.41, p=0.002), density of SOX2+ cells in the EGL (T, arrowheads n≥4/condition, Two-way ANOVA, F_(1,15)_=58.49, p<0.0001) and percentage of EdU+ SOX2+ cells within the SOX2+ cells in the BgL (U, arrows, n≥4/condition, Two-way ANOVA, F_(1,15)_=6.549, p=0.02). Scale bars: B-E: 500μm, F-I: 100μm, K-R: 50μm

We next asked whether the number of Bg or astrocytes generated was altered in P30 *Hopx*-*Ascl1* CKOs. Interestingly, there was a mild but significant increase in the density of Bg in IR *Hopx*-*Ascl1* CKOs compared to IR controls, but no differences between the other three groups. One possibility is that after irradiation the mutant BgL-NEPs generate Bg rather than GCPs (Figure S6L-P). In addition, the density of the astrocytes in the lobule WM in the IR *Hopx*-*Ascl1* CKOs was increased compared to nonIR mutants and IR controls (Figure S6Q-R). Finally, we quantified the density of ML INs, because we observed a population labeled with *Hopx*^*CreERT2*^ upon TM administration at P0. No significant changes in the densities of ML INs were observed after irradiation in the *Hopx*-*Ascl1* CKOs compared to their control littermates and their respective nonIR controls (Figure S6S-T). However, since the size of the mutant CB is significantly reduced, by extrapolation the total number of ML INs is reduced in IR mutants.

In order to understand why the regeneration is impaired in the *Hopx-Ascl1* CKOs, we analyzed nonIR and IR cerebella at P7, and determined whether proliferation of NEPs and/or their migration to the EGL were affected. Quantification of the density of ASCL1+ cells in the BgL in P7 IR and nonIR cerebella confirmed loss of ASCL1-expressing transitory cells after IR and thus deletion of *Ascl1* in mutants (Figure 6K-N, S). Significantly, *Hopx*-*Ascl1* CKOs had a lower density of SOX2+ cells in the EGL (BgL-derived cells that had migrated to the EGL) after injury compared to their control IR littermates (Figure 6O-R, T). A previous study suggested that proliferation genes are direct targets of ASCL1 in embryonic stem cell-derived neural stem cell cultures or the embryonic ventral brain (Castro et al., 2011). To test if loss of *Ascl1* in the *Hopx*-expressing NEPs impairs their proliferation, we injected EdU 1h prior to sacrificing P7 pups (Figure 6A) and assed the percentage of EdU+ SOX2+ cells in the BgL. A significant decrease in the percentage of EdU+ SOX2+ cells was detected in the BgL of IR *Hopx*-*Ascl1* CKOs compared to controls (Figure 6O-R, U). In summary, these results show that ASCL1 is required for the full reprograming of *Hopx*-expressing BgL-NEPs after depletion of GCPs, including efficient proliferation and migration to the EGL, and as a consequence of impaired replenishment of GCPs the cerebella of P30 IR *Ascl1* CKOs are reduced in size compared to control IR mice.

## Discussion

The neonatal mouse CB has remarkable regenerative potential and cellular plasticity upon injury. Using scRNA-seq and GIFM, we defined two NEP subtypes and the transcriptional signatures of their 5 subpopulations during homeostasis, and also identified a key new transitory cellular state that is necessary for adaptive reprogramming of BgL-NEPs into GCPs following EGL injury. Our results reveal that lineage propensity of NEPs (astroglial vs neural) is the primary molecular factor that differentiates NEP subtypes. At steady state *Hopx*-expressing NEPs are proliferative, primarily gliogenic and found in both the BgL and WM, whereas the *Ascl1*-expressing NEPs are proliferative but restricted to the lobule WM and dedicated to making ML INs at P5. Importantly, we discovered that upon depletion of the GCPs *Hopx*-expressing gliogenic BgL-NEPs transition to a new state where proneural VZ-bHLH genes are activated prior to their migration to the EGL. Furthermore, upon injury, we not only confirmed a delay in production of INs by neurogenic NEPs using GIFM but uncovered that a subset of neurogenic-NEPs switch to producing astrocytes, including some that become ectopically located in the deep WM. Finally, given that ASCL1 is a pioneer transcription factor (Wapinski et al., 2017) and we discovered it is required for full regeneration of the GCPs, we propose that ASCL1 is involved in erasing the gliogenic differentiation program of the BgL-NEPs, thus allowing them to acquire a GCP identity (Figure 7).

**Figure 7.**
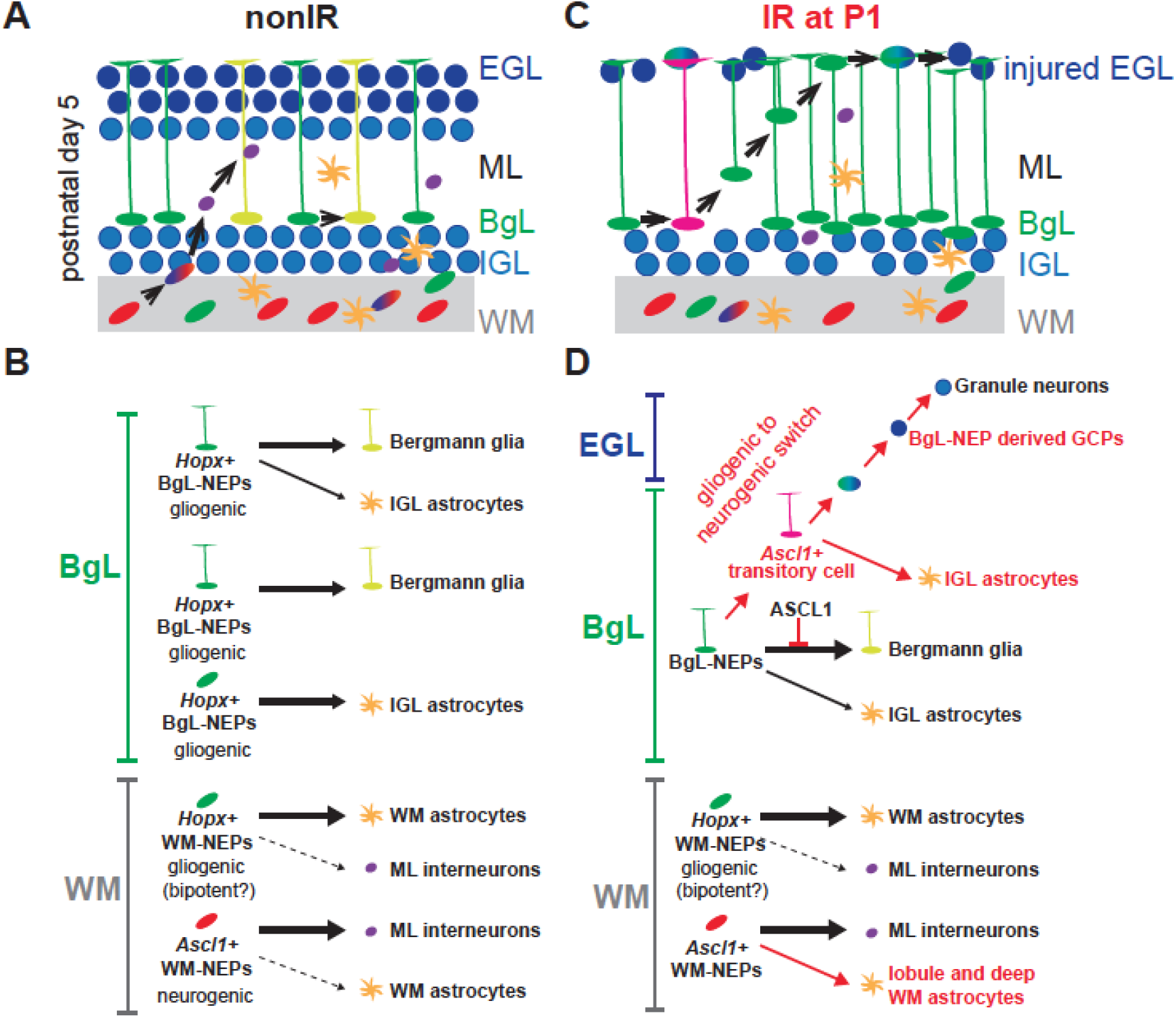
Schematic summary of the proposed NEP populations and lineages during homeostasis and repair. **A-B.** Based on our results and published clonal analyses (Cerrato et al., 2018; Parmigiani et al., 2015) we propose that within the two molecular NEP subtypes we identified, there are 5 NEP subpopulations in the cerebellar lobules at homeostasis. One molecular NEP subtypes is gliogenic (*Hopx*-expressing) and resides in the BgL and the WM and the other is neurogenic (*Ascl1*-expressing) and resides in the WM. scRNA-seq identified 3 subpopulations in the neurogenic NEPs that are omitted since they appear to be different stages of differentiation. **C-D.** Upon injury, a subset of *Hopx*-derived BgL-NEPs turn on a neurogenic transcriptional program, which suppresses their normal Bg fate and allows them to undergo adaptive reprograming to generate new GCPs. Upon injury, there is also an increase in astrocyte production from *Ascl1*-expressing NEPs. For simplicity, the two lineage dedicated BgL-NEP populations are omitted.

An important question is whether within the gliogenic-NEP subtype there are NEP subpopulations that are dedicated to making only Bg or astrocytes, and if so whether the latter are located in both the WM and BgL. Based on our scRNA-seq, there are two gliogenic-NEP subpopulations (nIR2 and nIR5) present at steady state and one is more related to astrocytes and the other to Bg. Whereas Bg only are located in the BgL, *Hopx*-derived astrocytes are located in the IGL and lobule WM, which raises the question of whether both *Hopx*+ BgL-NEPs and/or WM-NEPs generate astrocytes. For example, *Hopx-*BgL NEPs might generate IGL astrocytes and WM *Hopx*-NEPs generate the WM astrocytes. Elegant clonal labeling of apparently only BgL-NEPs using *Glast*^*CreER*/+^; *R26*^*Confetti*/+^ mice and application of Tm to the surface of the P6 CB *in vivo* found that the majority of clones had both Bg and astrocytes located in the IGL (~55%), although a large proportion were dedicated to making only Bg (~39%) and the rest to making only IGL astrocytes (Cerrato et al., 2018). This result indicates that WM astrocytes must be generated from WM-NEPs, likely the *Hopx*-expressing WM-NEPs we identified. Indeed, when WM progenitors were fluorescently labelled directly by injection of virus, some P3-P5 WM-NEPs gave rise to WM astrocytes (Parmigiani et al., 2015). Furthermore, our comparison of labeling before irradiation in IR and nonIR *Hopx-TdT* animals revealed preferential labeling of the BgL-NEPs that become *Ascl1*+ after injury, providing evidence that that not all BgL-NEPs are eqivalent in their extent of plasticity. Thus, molecular and lineage analysis shows that WM astrocytes and Bg have distinct progenitors, whereas IGL astrocytes might come from both a bipotent progenitor and lineage restricted progenitor in the BgL.

At steady state, *Ascl1* is expressed in all three neurogenic-NEP clusters, and our GIFM showed that *Ascl1*+ cells are exclusively in the lobule WM and give rise to ML INs at P5, in agreement with previous findings at other postnatal ages (Brown et al., 2019; Sudarov et al., 2011). Our differential expression analysis of the neurogenic clusters (nIR1, 3 and 4) showed they are very similar and suggests the main differentially expressed genes are related to cell cycle or *Cfp* level. One interpretation is that the three neurogenic WM-NEP subpopulations represent progressive differentiation states. Interestingly, we found that *Hopx-*expressing NEPs give rise to a small number of INs at P5 and the proportion and the number is higher when Tm is injected at P0, indicating there could be a bipotent *Hopx*+ WM-NEP that produces INs. Further clonal analysis is required to resolve whether gliogenic WM-NEPs can give rise to both INs and WM astrocytes.

We showed that induction of the proneural VZ-bHLH transcription factor ASCL1 in BgL-NEPs is a key step in their adaptive reprograming upon GCP death. Of importance, this *Ascl1*-driven gliogenic to neurogenic switch occurs without outside intervention such as the forced ectopic expression of proneural transcription factors required in other CNS regions (Grande et al., 2013; Heinrich et al., 2014; Zamboni et al., 2020). Further investigation is needed to identify the injury induced signal(s) that activate *Ascl1* expression. Since upregulation of *Ascl1* peaks 3-4 days after injury, the critical events required to produce the *Ascl1*+ transitory cell state must occur befor P5. Interestingly, in another study we showed that upon depletion of Purkinje cells in the newborn CB, although new Purkinje cells are generated, BgL-NEPs do not undergo adaptive reprogramming to produce them (Bayin et al., 2018). Thus, the type of injury/cell type killed is instrumental in determining the downstream regenerative response of NEPs.

We previously showed that at the same time that the BgL-NEPs are generating new GCPs after irradiation, the lobule WM-NEPs reduce their proliferation and production of INs and to a lesser extent astrocytes (Wojcinski et al., 2017). Our finding that expression of the WM-NEP marker *Igfbp5* is reduced in WM-NEPs provides molecular evidence for such a response to EGL injury. Importantly, a surprising observation was that after injury *Ascl1-*expressing NEPs produce a significant number of astrocytes located not only in the IGL and lobule WM, but some that migrate ectopically to the deep WM. Although the transitory *Ascl1*+ BgL-NEPs might be the source of the IGL astrocytes, it is likely that *Ascl1*+ WM-NEPs in the lobules have sufficient plasticity to switch to producing astrocytes in the lobules and deep WM upon injury. The reason for this neural to glial fate switch after injury and whether the same NEPs retain their neurogenic potential (i.e. are bipotent) or whether a distinct subpopulation of the *Ascl1*+ WM-NEPs become gliogenic remains to be explored.

In conclusion, our studies provide molecular insights into the make up of NEP subtypes and the endogenous permissive mechanisms that support a glial to neural switch crucial for enhancing repair after brain injury. A new transitory BgL-NEP population was identidied and we demonstrate that ASCL1, which normally drives cerebellar IN production (Sudarov et al., 2011), is required for adaptive reprograming of BgL-NEPs to produce excitatory granule cells upon EGL injury. The mechanisms and cellular states we identified may well have broader implications for gliogenic progenitors in other regions of the brain, and their potential responses to injury.

## Methods

### Animals

All the mouse experiments were performed according to protocols approved by the Memorial Sloan Kettering Cancer Center’s Institutional Animal Care and Use Committee (IACUC, protocol no 07-01-001). Animals were housed on a 12-hour light/dark cycle and given access to food and water ad libitum.

The following mouse lines were used: *Nes-Cfp* (Mignone et al., 2004; Wojcinski et al., 2017), *Hopx*^*CreERT2*^ (Takeda et al., 2011), *Ascl1*^*CreERT2*^ (Kim et al., 2011; Sudarov et al., 2011), *Ascl1*^*fl/fl*^ (Pacary et al., 2011; Sudarov et al., 2011) *Rosa26*^*lox-STOP-loxTdTomato*^ (ai14, Stock no: 007909, The Jackson Laboratories)(Madisen et al., 2015). Animals were maintained on an outbred Swiss Webster background. Both sexes were used for analyses and experimenters were blinded for genotypes whenever possible.

Tamoxifen (200ug/g, Sigma) was injected subcutaneously at P0 or at P5. EdU (50ug/g) was injected intraperitoneally at P7.

Analysis of *Ascl1-TdT* and *Hopx-TdT* P30 brains that were not injected with Tm showed no ectopic TdT+ cells (Figure S2K-L).

### Irradiation

P1 pups were anesthetized by hypothermia and a single dose of 4 Gy gamma-irradiation was provided using an X-RAD 225Cx (Precision X-ray) microirradiator in the Small Animal Imaging Core Facility at Memorial Sloan Kettering Cancer Center. The CB region was targeted using a collimator with 5 mm diameter.

### Tissue Preparation and Histological analysis

For P5 and younger animals, brains were dissected and drop fixed in 4% PFA for 24-48 hours at 4°C. Animals older than P5 were systemically perfused with ice-cold PBS followed by 4% PFA, following anesthesia with a Ketamine (100mg/kg) and Xylaxine (10mg/kg) cocktail. After dissection brains were fixed for an additional 24-48 hours in 4% PFA. Fixed brains were switched to 30% Sucrose in PBS. Once they sunk, brains were embedded in OCT (Tissue-Tek) for cryosectioning. 14um-thick sections were obtained using a cryostat (Leica, CM3050S) and stored at −20°C.

Haematoxylene and Eosin (H&E) staining was performed for cerebellar area (size) measurements and assessment of the cerebellar cytoarchitecture.

Immunofluorescent (IF) analysis was performed on cyrosections. Slides were allowed to warm to room temperature (RT) and washed once with PBS. 1 hour blocking was performed at RT using 5% Bovine Serum Albumin (BSA, Sigma) in PBS-T (PBS with 0.1% Triton-X). For ASCL1 IF analysis, Mouse on Mouse Blocking reagent (Vector Labs) was applied for 2 hours at RT to reduce the background. Slides were incubated with primary antibodies diluted in blocking solution at 4°C overnight (Table S6). Slides were then washed with PBS-T (3 × 5 min) and incubated with fluorophore-conjugated secondary antibodies (1:500 in blocking buffer, Invitrogen). Hoechst 33258 (Invitrogen) was used to label the nuclei and the slides were mounted with Fluoro-Gel mounting media (Electron Microscopy Sciences). To detect EdU, a Click-it EdU assay with Sulfo-Cyanine5 azide (Lumiprobe corporation, A3330) was used.

### RNA *in situ* hybridization

Specimen treatment and hybridization were performed as previously described (Blaess et al., 2011). Templates for the probes were *in vitro* transcribed either from PCR-amplified cDNAs obtained from neonatal cerebella extracts or synthesized DNA (GeneScript). An *Ascl1* probe was generated from a plasmid as previously described (Guillemot and Joyner, 1993). Sequences of the probes are shown in Table S7.

### Slice cultures and live imaging

P7 cerebella were dissected in ice-cold 1x HBSS (Gibco) and embedded in 4% low melting point agarose. 250 μm thick sagittal slices were obtained using a vibratome (Leica) and immediately placed on Millicell (Millipore) tissue culture inserts over Neurobasal media supplemented with 1x B27 and 1x N2 supplements (Gibco), 2mM L-glutamine and allowed to equilibrate for 30 minutes at 37°C in 5% CO_2_ prior to imaging. Image stacks were obtained every 3-4 minutes for 6-8 hours using a LSM880 (Leica) with an environmental chamber (37°C with 5% CO_2_). Movies were processed using ZEN (Leica) and ImageJ (NIH) software.

### Image acquisition and analysis

Images were collected with a DM6000 Leica microscope or Zeiss LSM 880 confocal microscope. Images were processed using ImageJ Software (NIH).

Three near midline sections for each animal were quantified and averaged for all the analyses shown. Quantification of the number of cells was performed on lobules 3 and 4/5 at P7 and on lobule 3 at P30. Boundaries of the lobules to be quantified were decided based on straight lines drawn from the bases of the fissures. Cell densities were calculated by dividing the number of cells by the area of the lobule(s) quantified. CB area was measured on H&E stained slides from 3 near midline sections and the values were averaged. The number of animals that is used in each quantification is denoted in the figure legends and wherever summary statistics are presented.

### Single cell sequencing and data analysis

#### Sample preparation

Four male P5 nonIR and IR cerebella were dissected into ice-cold 1x HBSS (Gibco) and were pooled for downstream analysis. All the steps were performed on ice when possible. Cerebella were minced with a clean blade and then dissociated in Accutase (Innovative Cell Tech.) at 37°C for 10-15 minutes. Following dissociation, accutase was washed out using Neural stem cell media (Neurobasal, supplemented with N2, B27 and non essentital amino acids, Gibco). Following filtering through a cell strainer and trituration in media to single cells, cells were layered over a 5mL density gradient (albumin-ovomucoid inhibitor solution, Worthington) and centrifuged at 70g for 6 minutes to remove debris. The cells were briefly treated with 1:2 1X red blood cell lysis buffer (Sigma) for 2 minutes at room temperature. Cells were washed twice (500g, 5 minutes at 4°C) and resuspended in 1X Tris-buffered Saline. Cells were stained with Calcein AM live stain dye (Fisher, 1:500) on ice for 30 minutes, and were passed through a 40 μm cell strainer to remove any cell clumps before loading onto a microwell device.

#### Single Cell library preparation and sequencing

On-chip reverse transcription, cDNA amplification and sequencing library preparation were performed as described previously (Mizrak et al., 2020; Mizrak et al., 2019; Yuan and Sims, 2016). Briefly, reverse transcription reactions were performed on DropSeq beads (Chemgenes, MACOSKO-2011-10 (V+)). The microwell devices were scanned during the RNA capture to check for lysis efficiency. Following PCR amplification, the libraries were prepared using Nextera XT kit. DNA purifications were performed using Ampure XP beads. cDNA and library amount and quality were assessed using Qubit and a Bioanalyzer (Agilent). High quality samples were sequenced on a NextSeq 500 using a High Output 75 cycle kit (Read 1: 21 Cycles, Index: 8 Cycles, Read 2: 63 cycles).

#### Single Cell RNA-sequencing Data Analysis

The sequencing reads were demultiplexed, aligned, and quantified as described previously (Mizrak et al., 2020). 12 nucleotide cell barcodes and 8 nucleotide unique molecular identifiers were extracted from Read 1, and trimmed Read 2 reads were aligned to a mouse genome (GRCm38, Gencode annotation vM10) using STAR v.2.5 (Dobin et al., 2013). *Cfp* sequence was added in the annotation and denoted as NestinCFP in the expression matrices.

Unsupervised clustering was carried out as described previously (Levitin et al., 2018; Mizrak et al., 2019). The clustering was performed on protein-coding genes only, including NESTINCFP. After computing cell-by-cell Spearman correlation matrix, k-nearest neighbors graph was constructed with k set to 30. The resulting graph was used as input for Louvain clustering with Phenograph (Levine et al., 2015). Cluster-specific genes were identified using a binomial-specificity test (Shekhar et al., 2016). Uniform Manifold Approximation and Projection (UMAP) was used for all data visualization (Becht et al., 2018).

Differentially expressed genes in nonIR compared to IR conditions were identified using Single-cell differential expression (SCDE) analysis (Kharchenko et al., 2014). The resulting gene lists were pre-ranked based on the effect size, and were inputted to GSEA (Subramanian et al., 2005) using the following parameters: xtools.gsea.GseaPreranked -scoring_scheme weighted –setmin 1 –setmax 1000 –nperm 1000.

To address co-expression and mutual exclusivity of genes detected in more than 25 cells, we calculated the marginal detection probabilities for each gene pair as described previously (Mizrak et al., 2020). The top 50 genes that were co-expressed with *Ascl1* in nonIR and IR cerebella were identified for downstream analysis. The co-expression values were plotted as a scatter plot to identify condition specific co-expressed genes. We projected the scRNA-seq profiles from each IR cluster into the UMAP embedding of the nonIR profiles as described previously (Szabo et al., 2019) using the *transform* function in Python implementation of UMAP (Becht et al., 2018). The code for this analysis is available at www.github.com/simslab/umap_projection.

We performed differential expression analysis between clusters within the IR or nonIR data as follows: First, we sub-sampled each pair of clusters to the same number of cells. Next, we sub-sampled the counts for each pair of clusters to the same average number of counts per cell, keeping only transcripts annotated as protein-coding. We normalized the resulting sub-sampled count matrices using the *computeSumFactors* function (Lun et al., 2016) in *scran* and then conducted differential expression analysis with the Mann-Whitney U-test as implemented in the *mannwhitneyu* function in SciPy and corrected the resulting p-values using the Benjamini-Hochberg method as implemented in the *multipletests* function in the Python module StatsModels.

### Statistical analysis

Prism (GraphPad) was used for all statistical analysis. Statistical tests performed in this study were Student’s two-tailed t-test, Two-way analysis of variance (ANOVA), followed by post hoc analysis with Tukey’s multiple comparison tests. p-values of the relevant post hoc analyses are shown in the figures and in the Table S8. The statistical significance cutoff was set at p<0.05 and the data is presented as mean ± standard deviation (SD) of the mean. F-statistics and p-values are stated in the figure legends and relevant post hoc comparisons are shown in the figures. n≥3 mice were used for each experiment and the sample size for each experiment is stated in the figure legends and the text.

## List of additional supplementary materials

**Movie 1**. Live imaging of the lobule 3 of a P7 nonIR *Ascl1-TdT* CB that was given Tm at P5. Outline shows EGL. Time lapse images are shown in Figure S5.

**Movie 2**. Live imaging of the lobule 3 of a P7 IR *Ascl1-TdT* CB that was IR at P1 and given Tm at P5. Outline shows EGL. Time lapse images are shown in Figure S5.

**Table S1**. Binomial test results showing the cell-type specific genes in the P5 nonIR and IR whole cerebella and in nonIR and IR NEPs (related to Figures 1, 3, S1, S3)

**Table S2**. Differential expression analysis amongst nonIR clusters (related to Figure 1)

**Table S3**. GSEA results of the differentially expressed genes between the nonIR and IR NEPs (related to Figure S3)

**Table S4**. Differential expression analysis amongst IR clusters (related to Figure 3)

**Table S5**. Top genes that are co-expressed with *Ascl1* in nonIR and IR cells (related to Figure 3)

**Table S6**. List of antibodies

**Table S7**. Sequences complementary to the probes used for RNA *in situ* hybridization.

**Table S8**. Summary of the statistics

## Acknowledgements

We thank past and present members of the Joyner laboratory for discussions and technical help. We are grateful to the MSKCC Small Animal Imaging and Mouse Genetics Core Facilities for technical services and support. A XRad 225Cx Microirradiator was purchased by support from a Shared Resources Grant from the MSKCC GeoffreyBeene Cancer Research Center. Finally, we thank the Sulzberger Columbia Genome Center, supported by a National Cancer Institute Cancer Center Support Grant (P30CA013696), for technical services for the scRNA-seq experiments. This work was supported by grants from the NIH to ALJ (NINDS R01NS092096, NIMH R37MH085726) and a National Cancer Institute Cancer Center Support Grant (P30 CA008748-48). NSB was supported by postdoctoral fellowships from NYSTEM (C32599GG) and NIH/NINDS (K99/R00 NS112605-01). PAS was supported by R01NS103473 from NIH/NINDS.

## Code Availability

Data analysis code used for scRNA-seq is available at https://github.com/simslab.

---

**Figure S1.**
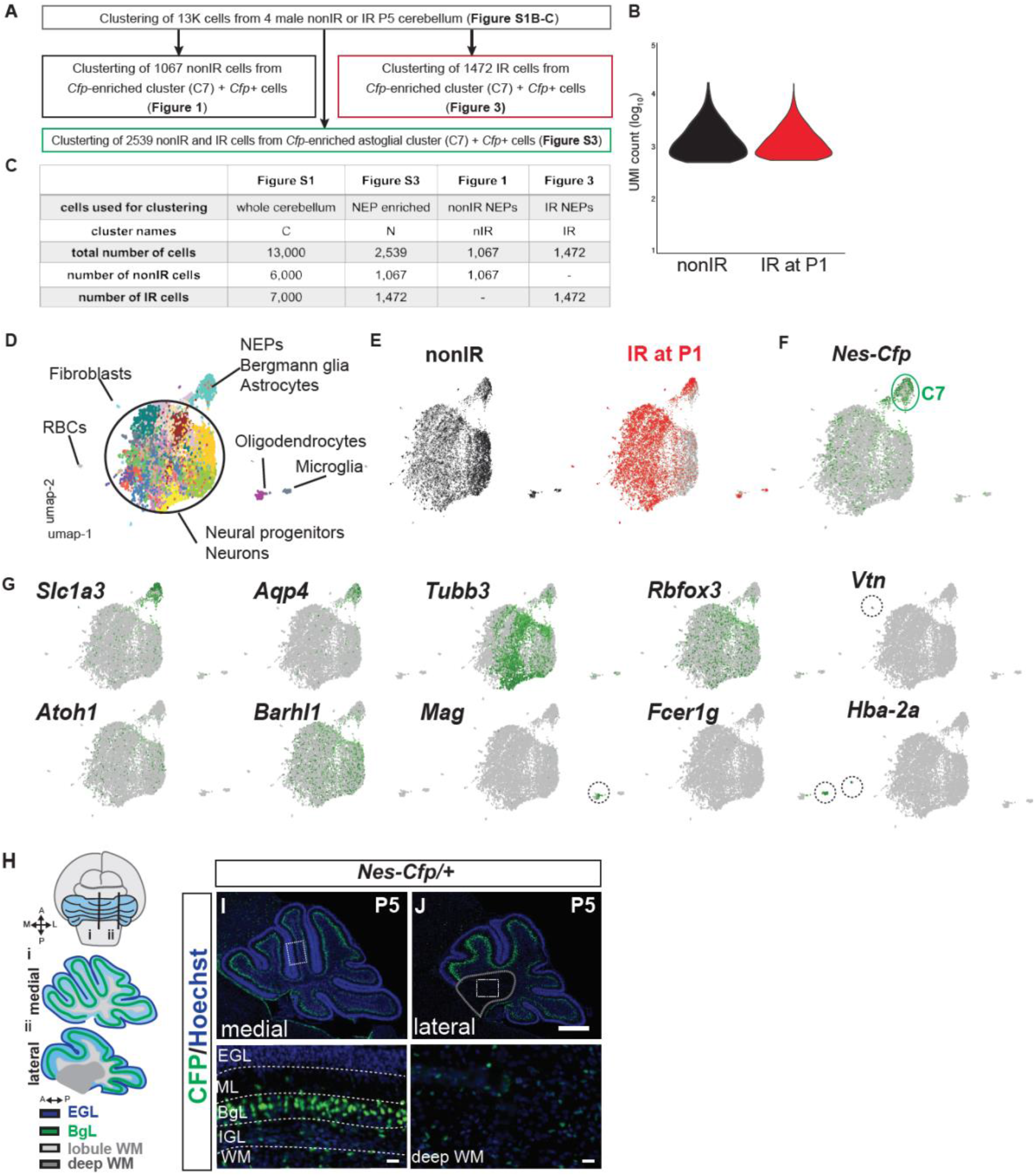
Summary of the scRNA-seq experiments and the initial clustering of the 13,000 cells from nonIR and IR cerebella at P5. **A**. Schematic explaining the clustering logic used for the scRNA-seq data analysis. **B**. Distribution of the UMI counts (log_10_) of the samples sequenced that passed quality control. **C.** Table summarizing the number of cells from nonIR and IR that were used for each analysis and the figures where each analysis is represented. **D-E**. UMAP visualization of the clustering of 13,000 cells divided by condition, nonIR (6,000) and IR (7,000) cells. 18 clusters were detected using Phenograph. **F.** Expression pattern of *Nes-Cfp* amongst the 13,000 cells. *Cfp* is significantly enriched in cluster C7 (circled). **G.** Expression patterns of markers for astrocytes/astroglia (*Slc1a3, Aqp4*), neurons l (*Tubb3, Rbfox3*), GCP/GCs (*Atoh1, Barhl1*), oligodendrocytes (*Mag*), microglia and macrophages (*Fcer1g*), red blood cells (*Hba-2a*) and fibroblasts (*Vtn*). **H.** Schematics showing the sagittal levels of the medial and lateral CB used for analysis. **I-J.** IF analysis of P5 *Nes-Cfp*/+ nonIR cerebella at medial (F) and lateral (G) levels showing the distinct localization of NEPs within the CB. Scale bars: 500μm, insets: 50μm

**Figure S2.**
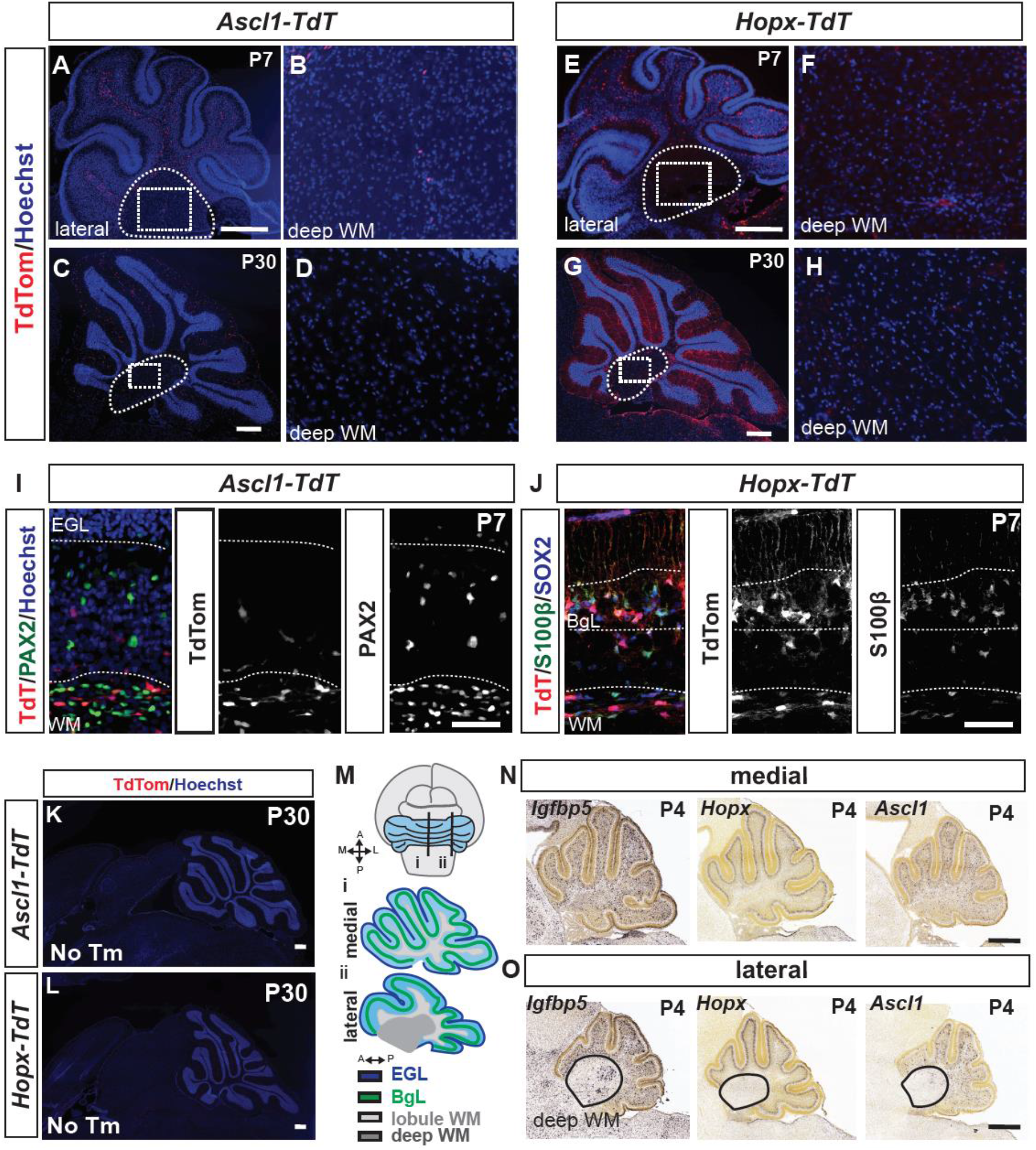
GIFM of *Hopx*+ and *Ascl1*+ NEPs and *RNA in situ* hybridization for cluster markers identified by scRNA-seq confirms NEP identities. **A-H**. Analysis of sections of the lateral CB of *Ascl1-TdT* and *Hopx-TdT* mice that houses the deep WM at P7 (A-B, E-F) and at P30 (C-D, G-H) shows no TdT+ cells in the deep WM. **I-J.** IF analysis at P7 of *Ascl1-TdT* and *Hopx-TdT* pups that were given Tm at P5 for the immature IN marker PAX2 (I) and Bg or BgL-NEP markers SOX2 (also all NEPs) and S100β, respectively (J). **K-L.** Analysis of P30 *Ascl1-TdT* and *Hopx-TdT* mice that were not given Tm show no ectopic labeling due to recombination. **E.** Schematics showing the level of sagittal levels in the medial and lateral CB used for analysis. **M-O.** Allen Brain Atlas P4 RNA *in situ* hybridization data showing the expression patterns of marker genes in clusters identified by scRNA-seq in the medial (N) and the lateral (O) CB. Scale bars: 500μm except I-J: 50μm

**Figure S3.**
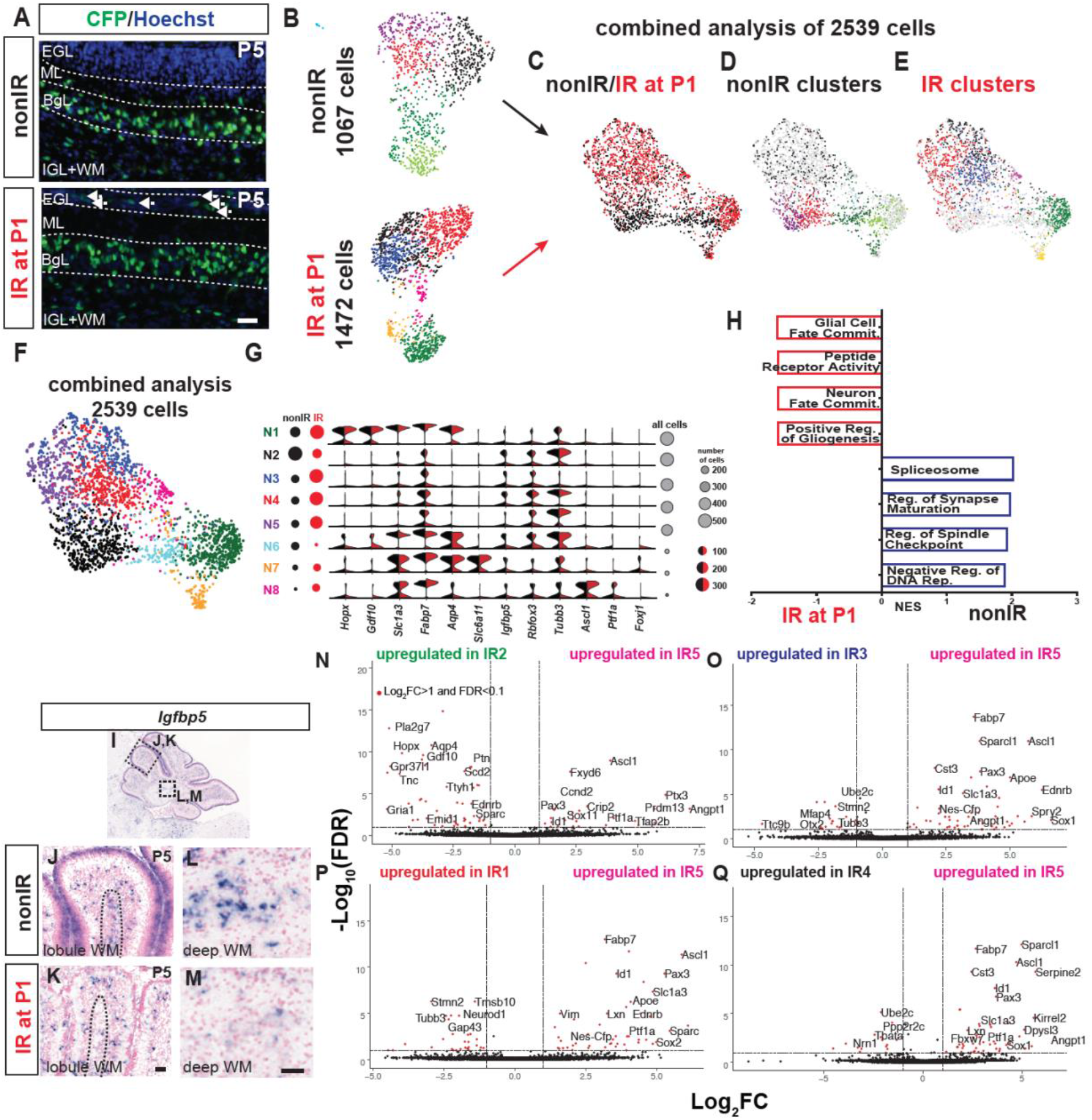
Clustering of nonIR and IR NEPs shows minimal batch affects and the pairwise comparisons of IR5 to other clusters highlights the distinct molecular signature of the *Ascl1*+ IR5 cells. **A.** IF analysis of P5 *Nes-Cfp*/+ nonIR and IR cerebella at P5**. B-E.** UMAP visualization of the clustering of 2,539 cells (1,067 nonIR and 1,472 IR cells) showing no batch affects in nonIR and IR samples. F-G. **H.** Clusters obtained from the analysis of 2,539 nonIR and IR NEPs and violin plots showing the log_10_CPM expression levels of canonical lineage genes and some of the top significantly enriched genes in the 8 clusters obtained in F. Circles show the total number of cells (black) or the nonIR (blue) or IR (red) number of cells within each cluster. **H.** GSEA results of differentially expressed genes shows the processes upregulated in nonIR vs. IR NEPs (NES>1.5, FDR<0.05). **I-M.** RNA *in situ* analysis of P5 nonIR and IR brains for the WM-NEP marker (lobule + deep WM, *Igfbp5*) identified by scRNA-seq. **N-Q.** Volcano plots showing the differentially expressed genes obtained from the pairwise comparisons of IR2 vs. IR5 (N), IR3 vs. IR5 (O), IR1 vs IR5 (P) and IR4 vs. IR5 (Q). Red dots show genes that are differentially expressed (fold change (FC)>Log_2_1 and false-discovery rate (FDR)<0.1). NES: normalized enrichment score. Scale bars: 50μm

**Figure S4.**
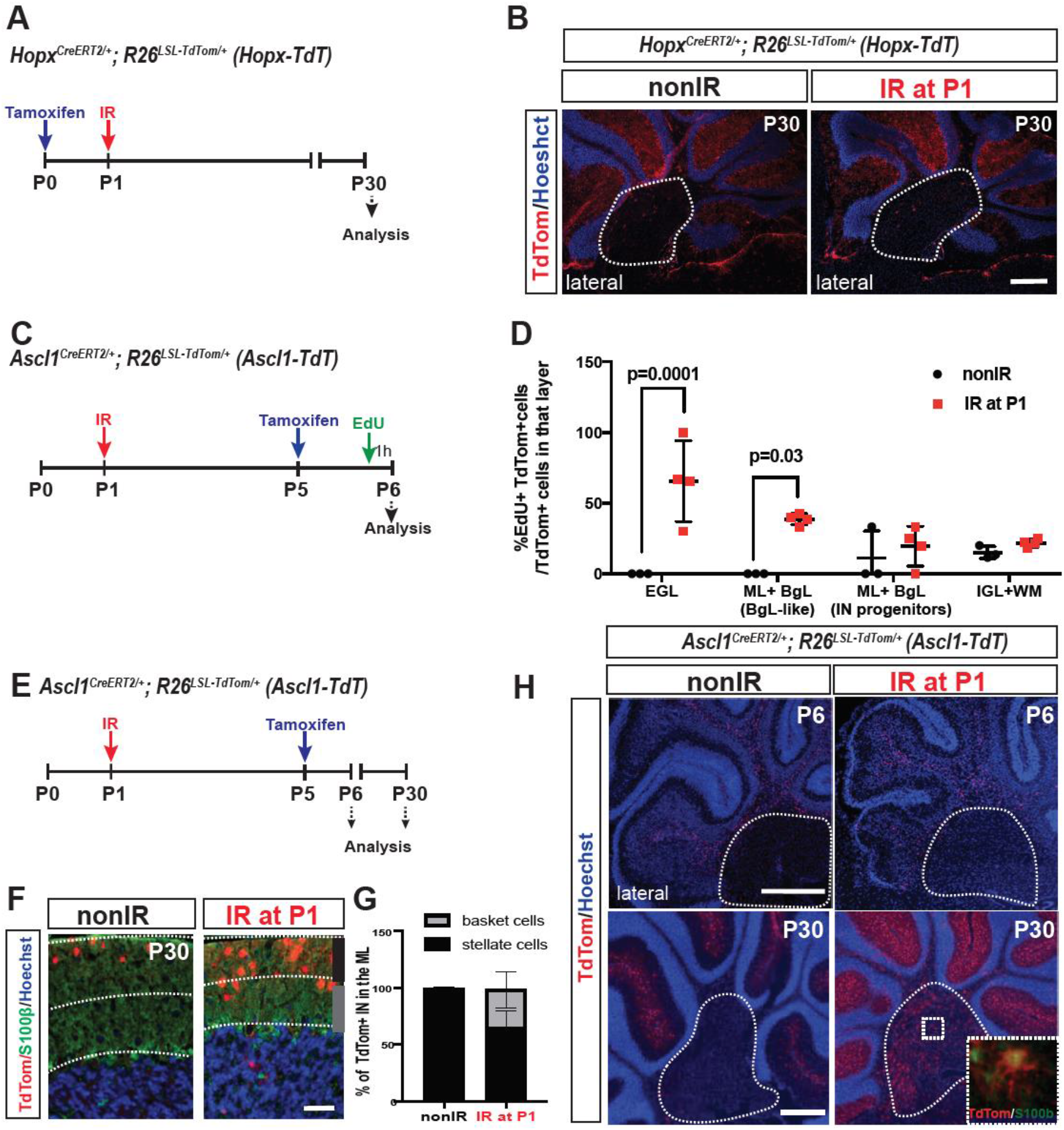
GIFM of *Hopx*+ gliogenic- and *Ascl1*+ neurogenic-NEPs in nonIR and IR brains shows changes in the lineages of the NEP subtypes. **A, C, E.** Schematics explaining the timelines of the experiments. **B.** Analysis of *Hopx-TdT* P30 nonIR and IR mice at P1 shows no TdT+ cells in the deep WM of the lateral CB. **D.** Quantification of the percentages of EdU+ TdT+ cells per all TdT+ cells in each layer in *Ascl1-TdT* animals at P6 in nonIR and IR P1 pups shows that the neurogenic-NEPs at steady state and the TdT+ BgL-NEPs and the TdT+ cells in the EGL cells are proliferative upon irradiation (n=3/condition, Two-way ANOVA, F_(3,20)_=31.28, p<0.0001). **F-G.** IF analysis of the TdT+ cells in the ML of the P30 nonIR and IR *Ascl1-TdT* animals (F) and the quantification of the proportions of Basket neurons (early born, lower ML) and Stellate neurons (later born, upper ML) shows a delay in the production of ML INs in the mice irradiated at P1. **H.** IF analysis of the lateral CB of nonIR and IR at P1 *Ascl1-TdT* animals shows TdT+ astrocytes in the deep WM (S100β+) at P30 but not at P6, suggesting the cells migrate from the lobule WM. Scale bars: 500μm, except for F: 50μm

**Figure S5.**
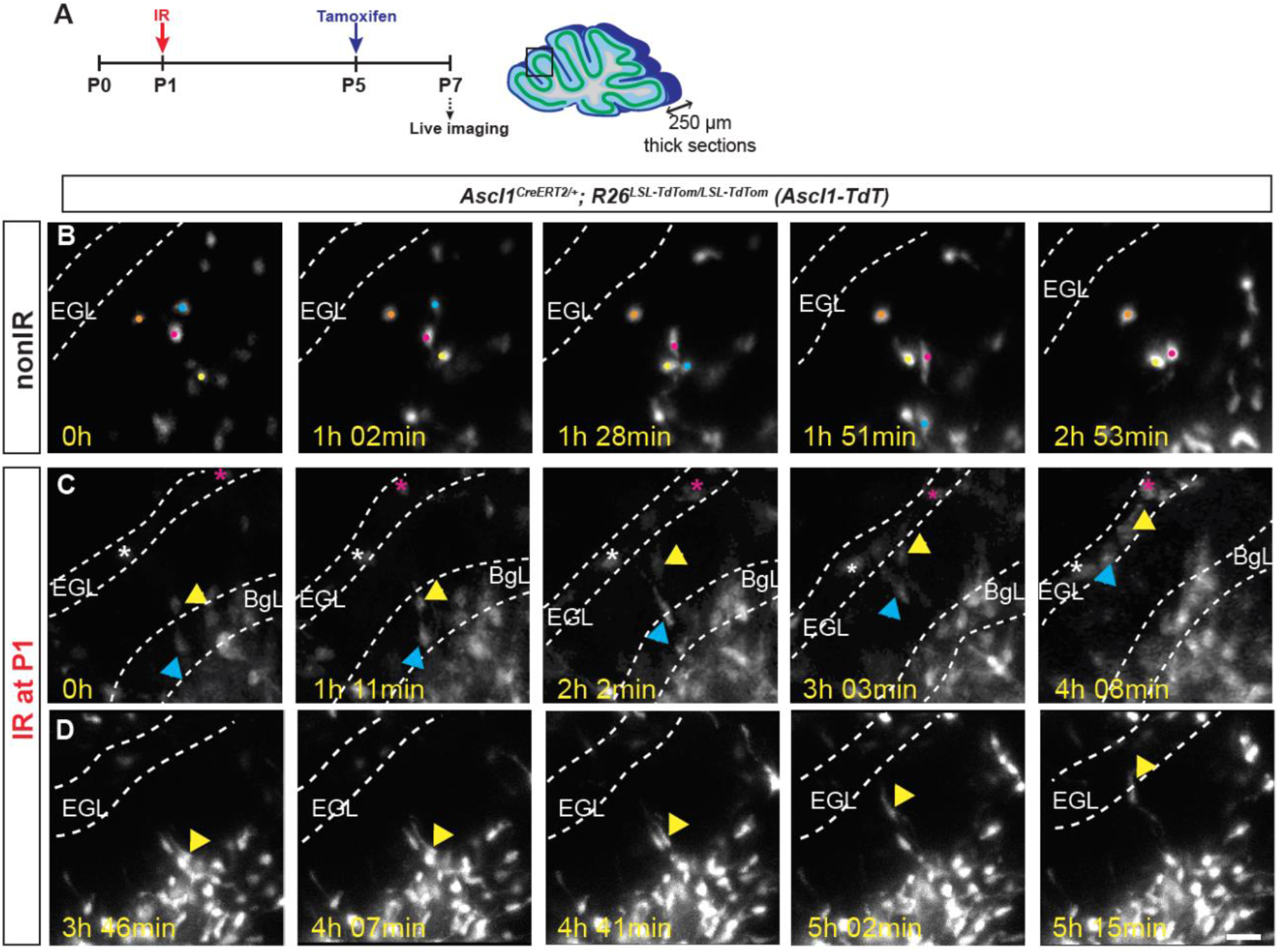
Time lapse images showing *Ascl1-TdT*+ BgL-NEPs migrating to EGL after IR. **A**. Schematic of the experimental plan. **B-D**. Examples of time-lapse live images of TdT+ cells from cerebellar sections obtained from P7 nonIR (B) or IR (C-D) *Ascl1-TdT* pups. Scale bars: 25μm

**Figure S6.**
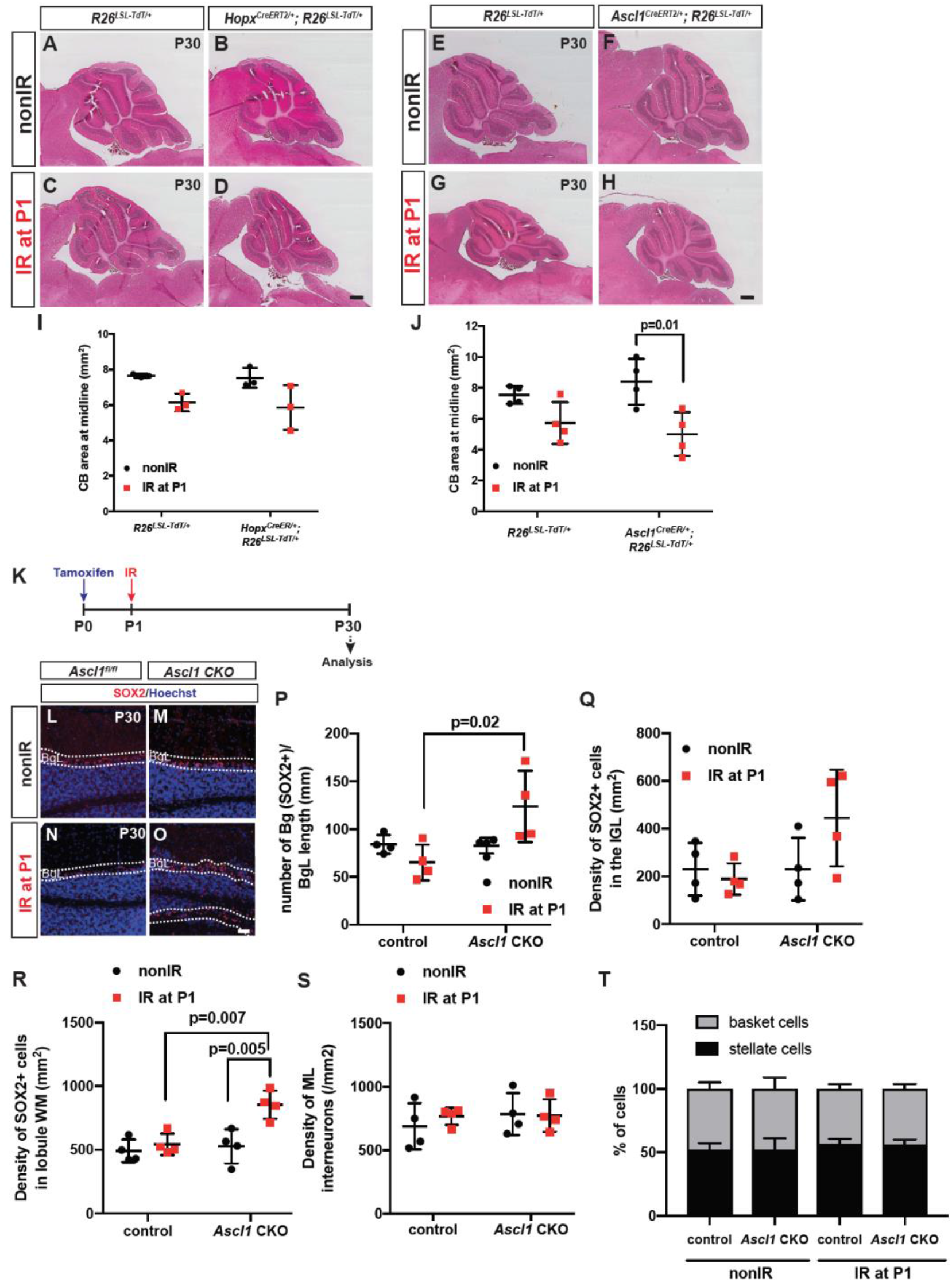
Loss of *Ascl1* in *Hopx*-NEPs increases Bg and astrocyte production upon irradiation but does not alter ML IN density. **A-J.** H&E analysis and the quantification of the cerebellar area at the midline of nonIR and IR, Cre+ and Cre-littermates of *Hopx-TdT* (A-D, I, n=3/condition, Two-way ANOVA, F_(1,4)_=13.41, p=0.02) and *Ascl1-TdT* (E-H, J, n=4/condition, Two-way ANOVA, F_(1,12)_=17.26, p=0.001) animals at P30. **K.** Experimental plan. **L-M.** IF analysis of control and *Ascl1* CKO nonIR and IR animals at P30 for SOX2 (Bg and astrocytes). **P-R.** Quantification of the density of SOX2+ Bg (P, n=4/condition, Two-way ANOVA, F_(1,12)_=6.878, p=0.02), and density of astrocytes in the IGL (Q, n=4/condition, Two-way ANOVA, p=0.086), and lobule WM (R, n=4/condition, Two-way ANOVA, F_(1,12)_=10.56, p=0.007). **S-T.** Quantification of ML IN density (S, n=4/condition, Two-way ANOVA, p=0.4) and the distribution of Basket vs. Stellate INs (T, n=4/condition) in nonIR and IR *Ascl1* CKO animals and control littermates shows no differences. Scale bars: A-H: 500μm, L-O: 100μm

**Movie 1.** Live imaging of the lobule 3 of a P7 nonIR *Ascl1-TdT* CB that was given Tm at P5. Outline shows EGL. Time lapse images are shown in Figure S5.

**Movie 2.** Live imaging of the lobule 3 of a P7 IR *Ascl1-TdT* CB that was IR at P1 and given Tm at P5. Outline shows EGL. Time lapse images are shown in Figure S5.

**Table S1.** Binomial test results showing the cell-type specific genes in the P5 nonIR and IR whole cerebella and in nonIR and IR NEPs (related to Figures 1, 3, S1, S3)

**Table S2.** Differential expression analysis amongst nonIR clusters (related to Figure 1)

**Table S4.** Differential expression analysis amongst IR clusters (related to Figure 3)

**Table S5.** Top genes that are co-expressed with *Ascl1* in nonIR and IR cells (related to Figure 3)

**Table S6.** List of antibodies

| Target | Catalog Number | Company | Dilution |
| --- | --- | --- | --- |
| goat α-SOX2 | AF2018 | R&D System | 1/200 (adult)-<br>1/500 (pups) |
| rabbit α-PAX2 | 71600 | Invitrogen | 1/500 |
| rabbit α-PAX6 | AB2237 | Millipore | 1/500 |
| rat α-GFP (CFP) | 04404-84 | Nacalai Tesque | 1/1000 |
| mouse α-ASCL1 | 556604 | BD Pharmingen | 1/1000 |
| guinea pig α-PVALB | 195-004 | Synaptic Systems | 1/1000 |
| rabbit α-S100B | Z0311 | DAKO | 1/1000 |
| mouse α-NeuN | MAB3777 | Millipore | 1/1000 |

**Table S7.**
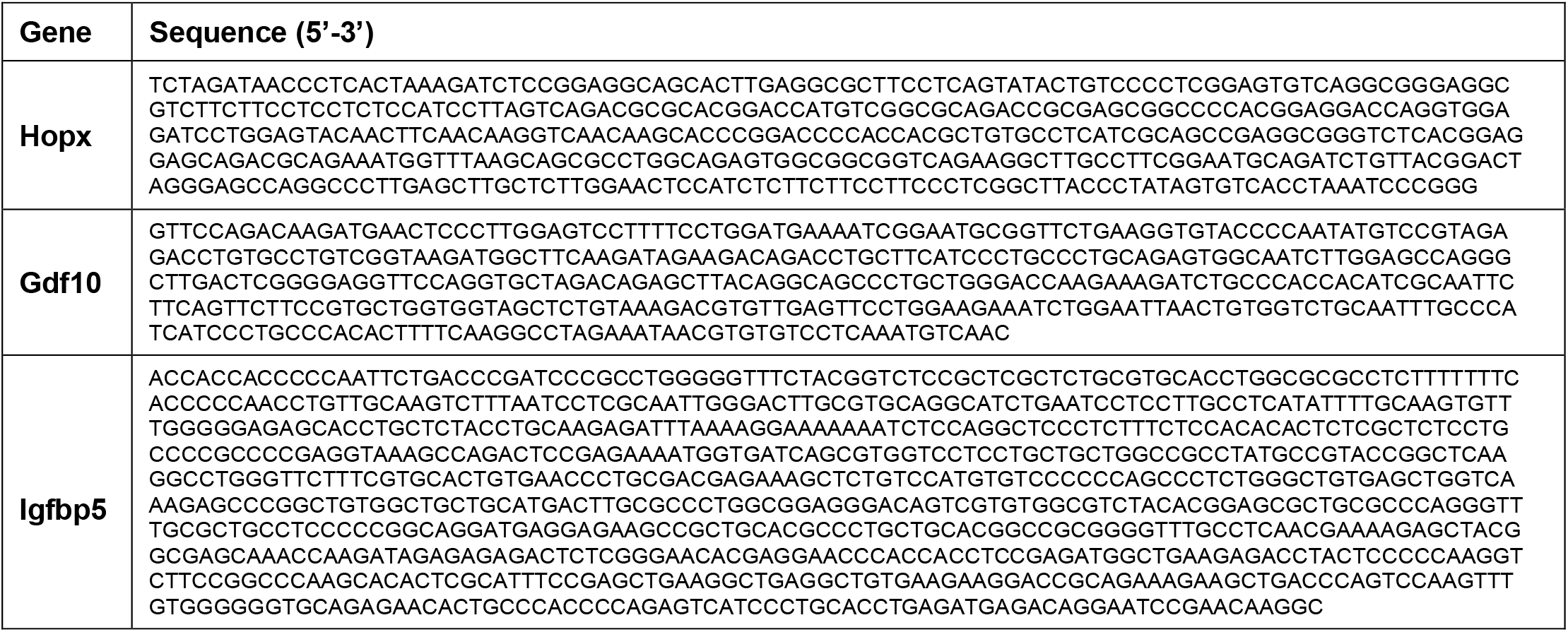
Sequences complementary to the probes used for RNA *in situ* hybridization.

**Table S8.** Summary of the statistics

| Figure | Test Performed | p-value | Multiple comparisons |  |
| --- | --- | --- | --- | --- |
| Figure 3N | Two-way ANOVA | $F_{(1,15)}=25.75$,<br>$p=0.0001$ | P5 | $p=0.001$ |
| Figure 4H | Two-tailed Student's t-test | $t=5.83$, $df=4$<br>$p=0.004$ | N/A | |
| Figure 4I | Two-way ANOVA | $F_{(1,8)}=7.928$,<br>$p=0.02$ | SOX2 IR vs. ASCL1 IR | $p=0.02$ |
| | | | ASCL1 nonIR vs. ASCL1 IR | $p=0.002$ |
| Figure 4P | Two-tailed Student's t-test | EGL: $t=5.5$,<br>$df=4$<br>$p=0.005$ | N/A | |
| Figure 4S | Two-tailed Student's t-test | GC: $t=3.513$,<br>$df=4$<br>$p=0.02$ | N/A | |
| Figure 4T | Two-way ANOVA | All layers:<br>$F_{(7,42)}=8.718$,<br>$p<0.0001$ | Granule neurons IGL nonIR vs. IGL IR | $p=0.004$ |
| Figure 5J | Two-way ANOVA | All layers:<br>$F_{(3,32)}=12.89$,<br>$p<0.0001$ | EGL 48h nonIR vs. IR | $p=0.008$ |
| | | | BgL 48h nonIR vs. IR | $p=0.007$ |
| Figure 5J | Two-way ANOVA | EGL only:<br>$F_{(1,8)}=5.94$,<br>$p=0.04$ | 24h IR vs. 48h IR | $p=0.03$ |
| | | | 48h nonIR vs. 48h IR | $p=0.003$ |
| Figure 5U | Two-way ANOVA | By layer and condition<br>$F_{(7,14)}=237.9$,<br>$p<0.0001$ | IN: ML nonIR vs. ML IR | $p<0.0001$ |
| | | | Granule neurons IGL nonIR vs. IGL IR | $p=0.0006$ |
| | | | Astrocyte WM nonIR vs. WM IR | $p=0.2$ |
| Figure 5U | Student's t-test for WM astrocytes | $t=4.966$,<br>$df=4$,<br>$p=0.007$ | N/A | |
| Figure 6J | Two-way ANOVA | by treatment:<br>$F_{(1,19)}=78.43$,<br>$p<0.0001$<br>by genotype:<br>$F_{(1,19)}=14.39$,<br>$p<0.001$ | control nonIR vs. IR | $p=0.0003$ |
| | | | mutant nonIR vs. IR | $p<0.0001$ |
| | | | control IR vs. mutant IR | $p=0.004$ |
| Figure 6S | Two-way ANOVA | by treatment:<br>$F_{(1,10)}=13.11$,<br>$p=0.005$<br>by genotype:<br>$F_{(1,10)}=16.41$,<br>$p=0.002$ | Control nonIR vs. IR | $p=0.004$ |
| | | | Control IR vs. mutant IR | $p=0.001$ |
| Figure 6T | Two-way ANOVA | by treatment:<br>$F_{(1,15)}=9.134$,<br>$p=0.009$<br>by genotype:<br>$F_{(1,15)}=58.49$,<br>$p<0.0001$ | control nonIR vs. IR | $p=0.00001$ |
| | | | mutant nonIR vs. IR | $p=0.03$ |
| Figure 6T | Two-way ANOVA | by treatment:<br>$F_{(1,15)}=9.134$,<br>$p=0.009$<br>by genotype:<br>$F_{(1,15)}=58.49$,<br>$p<0.0001$ | control IR vs. mutant IR | $p=0.002$ |
| | | | Control nonIR vs. IR | $p=0.00001$ |
| | | | Mutant nonIR vs. IR | $p=0.03$ |
| Figure 6U | Two-way ANOVA | by genotype:<br>$F_{(1,15)}=6.549$,<br>$p=0.02$ | control IR vs. mutant IR | $p=0.001$ |
| Figure S4D | Two-way ANOVA | treatment:<br>$F_{(3,20)}=31.28$,<br>$p<0.0001$ | EGL | $p=0.0001$ |
| | | | ML+BgL (BgL-like) | $p=0.03$ |
| Figure S6I | Two-way ANOVA | treatment:<br>$F_{(1,12)}=17.26$,<br>$p=0.0013$ | Post hoc multiple comparisons are not significant. | |
| Figure S6J | Two-way ANOVA | treatment:<br>$F_{(1,12)}=17.26$,<br>$p=0.0013$ | Mutant nonIR vs. IR | $p=0.01$ |
| Figure S6P | Two-way ANOVA | by genotype:<br>$F_{(1,12)}=6.878$ , p=0.02 | control IR vs. mutant IR | $p=0.02$ |
| Figure S6Q | Two-way ANOVA | by genotype:<br>$F_{(1,12)}=3.488$,<br>p=0.086 | N/A | |
| Figure S6R | Two-way ANOVA | by condition:<br>$F_{(1,12)}=12.49$<br>p=0.004<br>by genotype:<br>$F_{(1,12)}=10.56$<br>p=0.007 | mutant nonIR vs.<br>IR | p=0.005 |
| Figure S6Q | Two-way ANOVA | by genotype:<br>$F_{(1,12)}=3.488$,<br>p=0.086 | control IR vs.<br>mutant IR | p=0.007 |
| Figure S6S | Two-way ANOVA | by genotype:<br>$F_{(1,12)}=0.534$<br>p=0.4 | N/A | |

